# Transcriptomics-based screening identifies pharmacological inhibition of Hsp90 as a means to defer aging

**DOI:** 10.1101/468819

**Authors:** Georges E. Janssens, Xin-Xuan Lin, Lluís Millán-Ariño, Renée I. Seinstra, Nicholas Stroustrup, Ellen A. A. Nollen, Christian G. Riedel

## Abstract

Aging is a major risk factor for human morbidity and mortality. Thus, the identification of compounds that defer aging, also known as ‘geroprotectors’, could greatly improve our health and promote a longer life. Here we screened for geroprotectors, employing the power of human transcriptomics to predict biological age. We used age-stratified human tissue transcriptomes to generate machine-learning-based classifiers capable of distinguishing transcriptomes from young versus old individuals. Then we applied these classifiers to transcriptomes induced by 1300 different compounds in human cell lines and ranked these compounds by their ability to induce a ‘youthful’ transcriptional state. Besides known geroprotectors, several new candidate compounds emerged from this ranking. Testing these in the model organism *C. elegans*, we identified two Hsp90 inhibitors, Monorden and Tanespimycin, which substantially extended the animals’ lifespan and improved their health. Hsp90 inhibition specifically induces the expression of heat shock proteins, known to improve protein homeostasis. Consistently, Monorden treatment improved the survival of *C. elegans* under proteotoxic stress, and its lifespan benefits were fully dependent on the master regulator of the cytosolic unfolded protein response, the transcription factor HSF-1. Taken together, we present an innovative transcriptomics-based screening approach to discover aging-preventive compounds and highlight Hsp90 inhibitors as powerful geroprotectors that could be of great value, to target the aging process in humans.

## Introduction

Aging is widely accepted as a major risk factor for many diseases as well as mortality (Niccoli and Partridge, 2012). Thus, targeting the aging process directly by pharmacological means is increasingly viewed as a viable strategy to promote a healthier and longer life (Longo et al., 2015). Efforts are underway to explore these possibilities and to identify aging-preventive compounds, so-called ‘geroprotectors’. However, the list of candidates that are believed to confer such health and lifespan benefits to humans has remained very small (Barzilai et al., 2016; Kumar and Lombard, 2016). Even though screens covering tens of thousands of bio-active molecules have identified many drugs that extend lifespan in simple model organisms like yeast or worms (Lucanic et al., 2016; Petrascheck et al., 2007; Ye et al., 2014), validating their potential efficacy in humans is extremely time-consuming, limited in throughput, and restricted by ethical considerations. Thus, the sheer candidate numbers from such screens and their expected high frequency of a non-conserved effect in humans have discouraged their further evaluation (Kumar and Lombard, 2016).

One approach to improve the probability of identifying compounds that are effective in humans has been to screen for them in mammalian laboratory models such as mice, rats, or primates. But again, these studies are limited by significant costs, duration, and ethical considerations. Another approach has been to develop and employ screening methodologies directly in human that don’t require treatment of individuals, but instead limit themselves to compound screening in human cell-culture models and the computational interpretation of the resulting data. Although still in its early days, the latter approach has proven feasible, at least to identify dietary restriction mimetics (Calvert et al., 2016).

Here we tried to take such cell-culture-and computation-based approaches to a new level of sophistication: Recent studies had already demonstrated the power of human transcriptomes for biological age prediction (Peters et al., 2015; Sood et al., 2015; Yang et al., 2015). We decided to make use of this predictive power and to apply it to drug-induced transcriptional changes in human cell-culture models, to identify compounds that shift cells to a ‘younger’ state and thus may be geroprotective. We begin our screening efforts by using age-stratified human transcriptome datasets, covering over 50 tissues from donors between 20 and 69 years of age (a total of over 8000 transcriptomes from the Genotype-Tissue Expression ‘GTEx’ dataset (Lonsdale et al., 2013; Mele et al., 2015; The GTEx Consortium, 2015)), and build machine-learning algorithms (Kuhn et al., 2012; Liaw and Wiener, 2002) capable of distinguishing young versus old transcriptomes. Next, we apply these classifiers to transcriptomes resulting from drug responses to over 1300 unique compounds (provided by the Connectivity Map (CMap) (Lamb et al., 2006)) and rank drugs on a ‘geroprotective index’. Finally we validate our top hits in the model organism *Caenorhabditis elegans*. Applying this strategy, we identify several new geroprotector candidates, most notably the Hsp90 inhibitors Monorden (also known as Radicicol) and Tanespimycin. Focusing on these Hsp90 inhibitors, we determine the mechanism by which they exert their beneficial effects, namely by HSF-1-mediated improvement of protein homeostasis, and place our findings in the context of recent work that describes Hsp90 inhibitors as immuno-suppressants (Tukaj and Węgrzyn, 2016) and senolytics (Fuhrmann-Stroissnigg et al., 2017). Both of these function are absent in *C. elegans* but should even further potentiate the geroprotective benefits of Hsp90 inhibition when applied to humans.

## Results

### Construction of a core set of transcriptome-based age classifiers

Recent studies have demonstrated that biological age can be estimated by machine learning approaches applied to healthy tissue transcriptome datasets (Peters et al., 2015; Sood et al., 2015). We reasoned that applying such biological age-classifiers to drug-induced transcriptomes could reveal compounds with potentially youth-inducing properties (Fig. 1A). To test this, we required a dataset of transcriptomes from age-stratified donors. We thus turned to data made publically available by the GTEx consortium, which contained a diverse set of human tissue transcriptomes originating from donors of various ages and both genders (Lonsdale et al., 2013; The GTEx Consortium, 2015; Yang et al., 2015). This data has been used previously to study aging, showing its effectiveness in distinguishing age-related transcriptional signatures (Yang et al., 2015). Transcriptomes were downloaded from the GTEx consortium portal and preprocessed as previously described (Mele et al., 2015; Taskesen and Reinders, 2016; The GTEx Consortium, 2015), which resulted in a dataset of 8,555 transcriptomes from 51 tissues, both genders, and grouped into decade-sized age bins (ages 20-29, 30-39, 40-49, 50-59, and 60-69 (bins as provided by GTEx)) (Fig. 1B).

**Figure 1.**
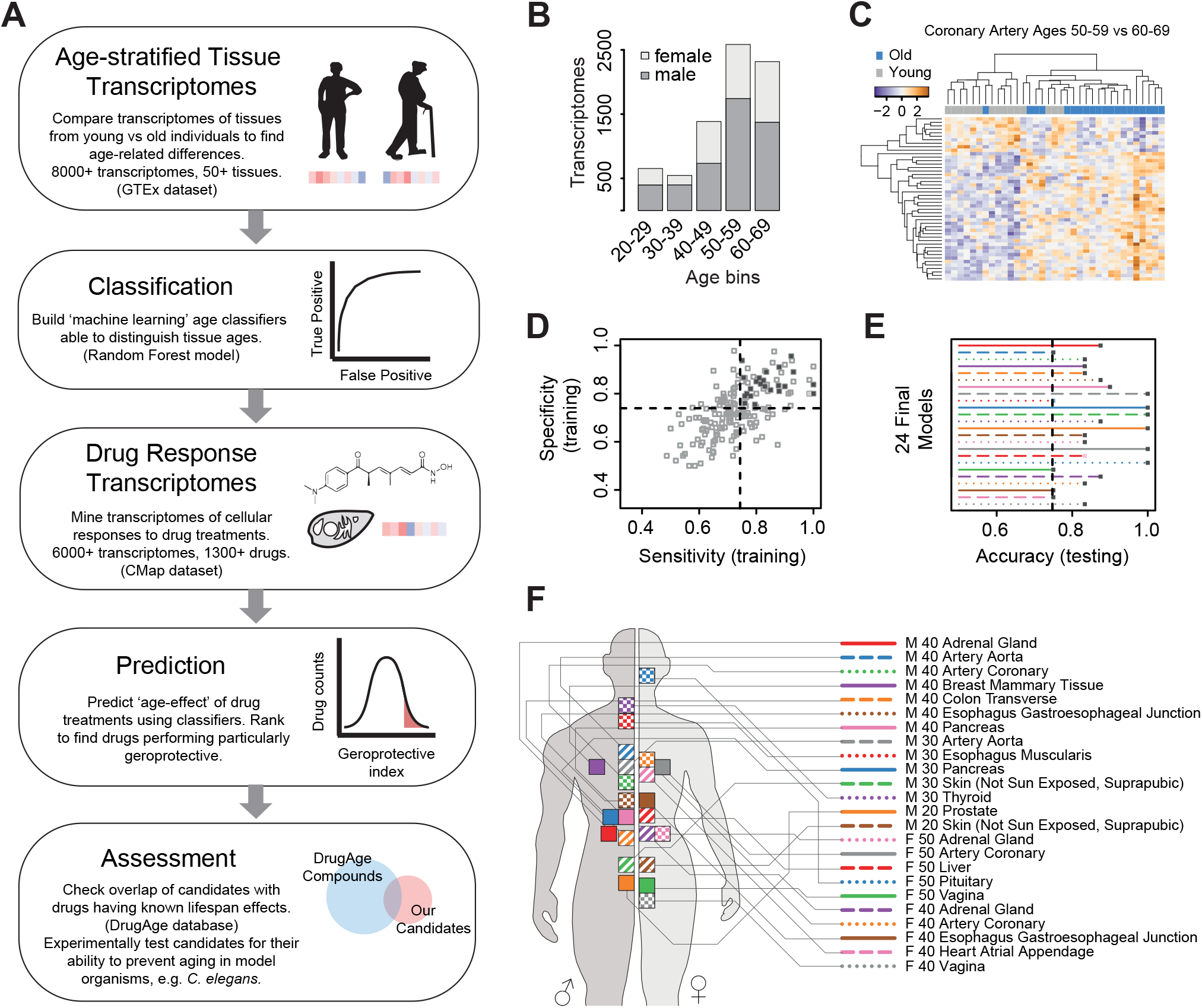
Study strategy and generation of transcriptome-based age classifiers. (A) General strategy for the discovery of geroprotective drugs. Transcriptomes of young versus old individuals (box 1) are used to generate age classifiers (box 2). These classifiers are then applied to drug-response transcriptomes (box 3), to identify compounds that change the transcriptome to a more ‘youthful’ state. A ranking is generated, prioritizing compounds that are most likely to be geroprotective (box 4). Finally, we assess the highest-ranked compounds against known geroprotectors to estimate the efficacy of our prioritization method (box 5). (B) The age and gender demographics of the GTEx data used in this study. (C) Representative heatmap of a binary ‘young’ (50-59) vs ‘old’ (60-69) comparison performed in a particular tissue (coronary artery). (D, E) Inclusion criteria for the final classification models, demanding sensitivity, specificity, and accuracy scores of above 0.75. (F) Distribution of the final 24 models among tissues and genders.

Next, we took a binary classification approach to train machine-learning models to be able to distinguish ‘young’ versus ‘old’ ages from transcriptional signatures. Tissue data was separated by gender and kept in the decade-sized age bins provided by the GTEx consortium. We defined being ‘old’ as the decade of 60-69 years (the oldest decade available in the GTEx data which matched our minimum sample number criteria) and made binary comparisons to any of the ‘younger’ decades. To minimize noise and limit the analysis to transcripts most likely to distinguish old from young samples, we filtered the datasets as follows: Of all possible binary comparisons, only those were made that contained at least 10 transcriptomes in both, the ‘young’ and ‘old’ datasets. The lowest 10% abundant transcripts were removed from the datasets to reduce noise. We further limited the analysis to genes which were differentially expressed between the two compared age bins, and finally we filtered to only include genes present in the Connectivity Map (Lamb et al., 2006), the transcriptional drug response dataset that we already mentioned in the introduction and will use in later stages of this study (Fig. 1C). Using these filtered datasets, we generated random forest models and tuned them in an automated, systematic manner (Kuhn et al., 2012; Liaw and Wiener, 2002) (Fig. 1, S1), resulting in 182 age-classification models with variable degrees of sensitivity, specificity, and accuracy (Fig. 1, Table S1). To generate a final core set of models, we applied cutoff criteria, including model sensitivity, specificity, and accuracy from the training and testing phases (Fig. 1D,E), ensuring that only the most effective classifiers remained. This resulted in 24 models covering both genders, various tissues, and various age comparisons (Fig. 1F, Table S2).

Our core set of 24 models was comprised of 1927 unique transcripts (Table S3). Although the majority of these genes had no known age-related functions, several prominent aging-related genes were present in this list and thus contributed to the age-classification: These included genes encoding the oxidative stress response proteins glutathione S-transferase pi 1 (GSTP1) and thioredoxin (TXN), where overexpression of the former was shown to increase lifespan in *C. elegans* (Ayyadevara et al., 2005) while overexpression of the latter increases lifespan in *Drosophila melanogaster* (Umeda-Kameyama et al., 2007) and mice (Mitsui et al., 2002), the insulin-like growth factor 1 receptor (IGF1R), orthologs of which affect lifespan in *C. elegans* (Kenyon et al., 1993), *Drosophila melanogaster* (Tatar et al., 2001), and mice (Holzenberger et al., 2003), the sirtuin SIRT1, also linked to aging in various model organisms (Cohen, 2004; Herranz et al., 2010; Kaeberlein et al., 1999; Rogina and Helfand, 2004; Tissenbaum and Guarente, 2001), and the mitochondrial uncoupling protein 2 (UCP2), overexpression of which increases lifespan in both *Drosophila melanogaster* (Fridell et al., 2005) and mice (Conti et al., 2006).

### Application of the age classifiers to rank compounds by their geroprotective potential

Having generated these 24 age-classification models, we then moved on to apply these models, in order to detect geroprotective compounds (Fig. 1A, steps 3-5). First, we required a suitable dataset of compound-induced transcriptional responses. For this we turned to the publicly available Connectivity Map (CMap), a resource consisting of over 6000 transcriptomes of various compound treatments performed on a selection of human cell lines (Lamb et al., 2006). In total, 1309 different compounds are covered by this dataset, including FDA approved medications but also a variety of other bioactive molecules. The CMap has successfully been used to find drugs affecting complex phenotypes, such as Celastrol for the treatment of diabetes (Liu et al., 2015) and Allantoin that acts as a caloric restriction mimetic (Calvert et al., 2016).

We reasoned that a cell line’s exposure to aging-preventive compounds would induce transcriptional changes classified as ‘young’ by our models (Fig. 1A). Before we could apply our models to the CMap data, we first had to account for the fact that the CMap data and our GTEx-derived models originate from different types of cells with distinct baseline gene expression profiles. Thus, for each age-classification model, we generated a prototypical ‘middle age’ transcriptome that was comprised of each gene’s average expression level between the ‘young’ and ‘old’ age group used to generate the model (see methods section for details). This resulted in transcriptomes that the models should not easily classify as either ‘young’ or ‘old’ (Fig. 2A). Next, we applied the fold gene expression changes of the drug responses in CMap to these prototypical ‘middle aged’ transcriptomes, and asked our models whether this would effectively shift the ‘middle aged’ transcriptome towards a ‘young’ classification and hence a more youthful state. Following this approach, each CMap perturbation entry was systematically applied to all of the 24 models’ specific ‘middle age’ transcriptomes (Fig. 2A), resulting in 24 different predictions for each of CMap’s more than 6000 entries (Table S4). To address the fact that the 1309 compounds in the CMap often have been evaluated at various doses, incubation times, or in different cell lines, and to give a drug the greatest possibility to show its geroprotective potential, we selected for each drug the CMap entry that gave the maximal probability of being young. This consolidated the dataset to 24 different predictions each across 1309 CMap drug entries. Furthermore, for better comparability between predictions from different models, we normalized model results to a maximal amplitude of 0.5, centered these probabilities around 0, and thereby created a ‘geroprotective index‘, in which a positive value signifies prediction of a rejuvenated transcriptome (Fig. 2B, Table S5).

**Figure 2.**
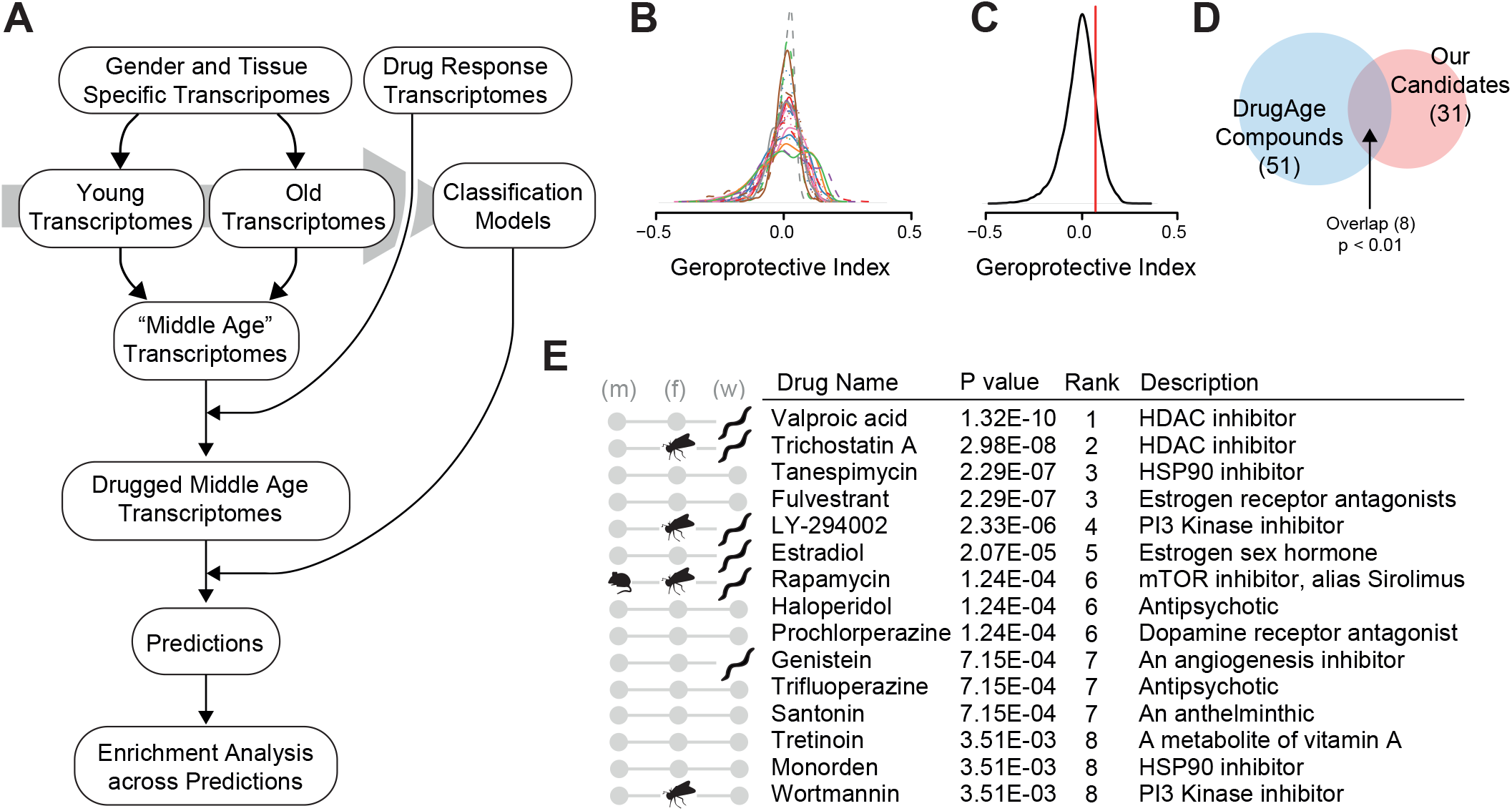
Drugs ranked by geroprotective index. (A) Pipeline for ranking CMap compounds by their geroprotective index. Tissues are separated into ‘young’ and ‘old’ groups, used to build models. ‘Middle age’ representative transcriptomes are generated by averaging between ‘young’ and ‘old’. CMap drug response fold changes are applied to the middle age transcriptomes to generate ‘drug-induced’ transcriptomes for each candidate drug. Age classifying models predict the ages of the ‘drug-induced’ transcriptomes. Enrichment scores are generated based on the 24 models’ separate predictions, to find drugs most often ranked as geroprotective. (B) Geroprotective index ranking of CMap compounds, for each of the 24 models (distribution of scores). The geroprotective index is a model’s prediction (the probability) whether a transcriptome should be classified as ‘young’. This probability was rescaled from −0.5 to 0.5, so that positive values would indicate a rejuvenating effect. (C) Consolidated geroprotective index scores (from (B)) of the CMap compounds (distribution, black line). The red line designates the mean absolute deviation, used as significance cutoff. (D) Overlap between our significant geroprotector candidates (31 total) and any CMap compounds that are listed as lifespan extending in the DrugAge database (51 total). (E) The top 15 geroprotective candidate compounds resulting from our method. Compounds published to increase lifespan in either mice (m), flies (f), or worms (w) are indicated. The candidates include Valproic acid, an HDAC inhibitor previously shown to increase lifespan in worms (Evason et al., 2008), Trichostatin A, another HDAC inhibitor previously shown to increase lifespan in both worms (Calvert et al., 2016) and flies (Tao et al., 2004), LY-294002, a PI3 Kinase inhibitor, shown to increase lifespan in both worms (Calvert et al., 2016) and flies (Moskalev and Shaposhnikov, 2010), Estradiol, shown to increase lifespan in worms (Ye et al., 2014), Rapamycin, also known as sirolimus, well known to extend lifespan in worms (Calvert et al., 2016), flies (Bjedov et al., 2010), and mice (Harrison et al., 2009), Genistein, an angiogenesis inhibitor, shown to extend lifepan in worms (Lee et al., 2015), and Wortmannin, another PI3 Kinase inhibitor, shown to extend lifespan in flies (Danilov et al., 2013).

To obtain a final evaluation for each drug, we assessed how frequently a drug was ranked as ‘geroprotective’ by the 24 models (Fig. 2C). We used the mean absolute deviation (Fig. 2C, red line) as a cutoff, summed the number of times (out of 24) a drug was above this cutoff, and performed enrichment tests in relation to the whole distribution of predictions. After correcting for multiple hypothesis testing, this resulted in a final significance-based ranking of drugs for their geroprotective potential (Table S6). We selected a corrected p value cutoff of 0.05 to form a candidate list of drugs for further consideration, which consisted of 31 compounds in total (Table S6; the top 15 compounds are shown also in Fig. 2E).

To assess the efficacy of our method, we explored what was known about these 31 top candidates. Comparing the overlap with the recently published DrugAge database (Barardo et al., 2017), a repository of compounds yielding lifespan effects in various model organisms, we found that of the over 400 unique drugs that significantly increase lifespan and thus are considered geroprotectors in DrugAge, 51 were present in CMap, and of these, 8 were in common with our list of 31 top candidates. This showed that our top candidates were significantly enriched for known geroprotective compounds (p < 0.01) (Fig. 2D, Table S7), confirming that our screening method worked well. Figure 2E shows the top 15 compounds from our candidate list, covering a wide range of mechanistic targets. These include the histone deacetlyase inhibitors Valproic Acid and Trichostatin A, the PI3 Kinase inhibitors LY-294002 and Wortmanin, as well as the TOR inhibitor Rapamycin (also known as Sirolimus), all of which have been shown to extend lifespan in various organisms (Bjedov et al., 2010; Calvert et al., 2016; Danilov et al., 2013; Evason et al., 2008; Harrison et al., 2009; Moskalev and Shaposhnikov, 2010; Tao et al., 2004) (Fig. 2E). Also several antidepressants were amongst our candidates, a class of drugs which has been shown to influence lifespan before (Petrascheck et al., 2007; Zarse and Ristow, 2008). Finally, in ranks 16 to 31 we found several drugs which have been suggested to protect from all-cause mortality in epidemiological studies in humans, in particular the diabetes therapeutic Metformin (Bannister et al., 2014), the rheumatoid arthritis therapeutic Methotrexate (Wasko et al., 2013), the acetylsalicylic acid (Aspirin) (Gum et al., 2001), as well as Clozapine, a drug used to treat serious mental illness (Hayes et al., 2015).

Taken together, our screening approach was able to successfully rank compounds by their geroprotective potential, revealing a top candidate list significantly enriched for known lifespan extending compounds. In addition, new geroprotector candidates were identified that should be interesting to investigate further.

For the scientific community, we provide an easy to use R script that uses our classifiers and can be used to make geroprotective predictions from novel drug-induced transcriptomes (Fig. S2 and supplemental R script data package).

### Validation of geroprotector candidates using *C. elegans*

To test for lifespan extending capabilities amongst our candidate compounds, we turned to the nematode *C. elegans* – a relatively short-lived model organism frequently used in aging research, including drug screening for geroprotectors (Calvert et al., 2016; Carretero et al., 2015; Ye et al., 2014). We selected 29 compounds for evaluation: These included 14 of the top 15 compounds from our candidate list (p-value cutoff: 3.6 x 10^-3^; Fig. 2E; one drug, Tanespemycin, was omitted due to target redundancy with Monorden and exceptional cost, although we retested it subsequently, as described below). Additionally, we included 15 compounds that we selected by hand (indicated in Table S6), irrespective of their ranking, but because we considered them promising geroprotector candidates due to their existing annotations, or in order to include drugs ranked poorly by our predictors (negative controls).

Compounds were evaluated at 50 µM (for the solvents used see Table S8), a concentration that was deemed optimal in the light of previously conducted *C. elegans*-based geroprotector screens (Calvert et al., 2016; Carretero et al., 2015; Ye et al., 2014). In order to obtain high resolution lifespan assays on this substantial panel of drugs, we used an automated imaging and analysis platform called the *C. elegans* ‘Lifespan Machine’ (Stroustrup et al., 2013). We applied the compounds to L4 stage larvae (animals near the end of their development), performed the survival assays, and eventually processed the data, making sure that each drug was tested using at least 50 worms. This resulted in a final list of 25 drugs giving high quality lifespan data, displaying a range of lifespan phenotypes from an almost 10% decrease to more than a 25% increase (Fig. 3A). Five compounds extended lifespan significantly and by more than 10%, namely Felbinac, Valproic Acid, LY-294002, Rapamycin, and Monorden – results that we reproduced by biological replicates (Fig. 3A, highlighted by red circles; Table S8; see Fig. 3B-D and Fig. S3A,B for individual survival curves). One compound (Tyrphostin AG-1478) that was significantly extending lifespan but did not meet our inclusion criteria for worm numbers in the initial screen, we later validated with larger numbers of worms (Fig. S3C, Table S8). Unfortunately, a few lifespan extension phenotypes previously published by other labs could not be confirmed by our assays, i.e. for Genistein, but this may be due to different experimental setups.

**Figure 3.**
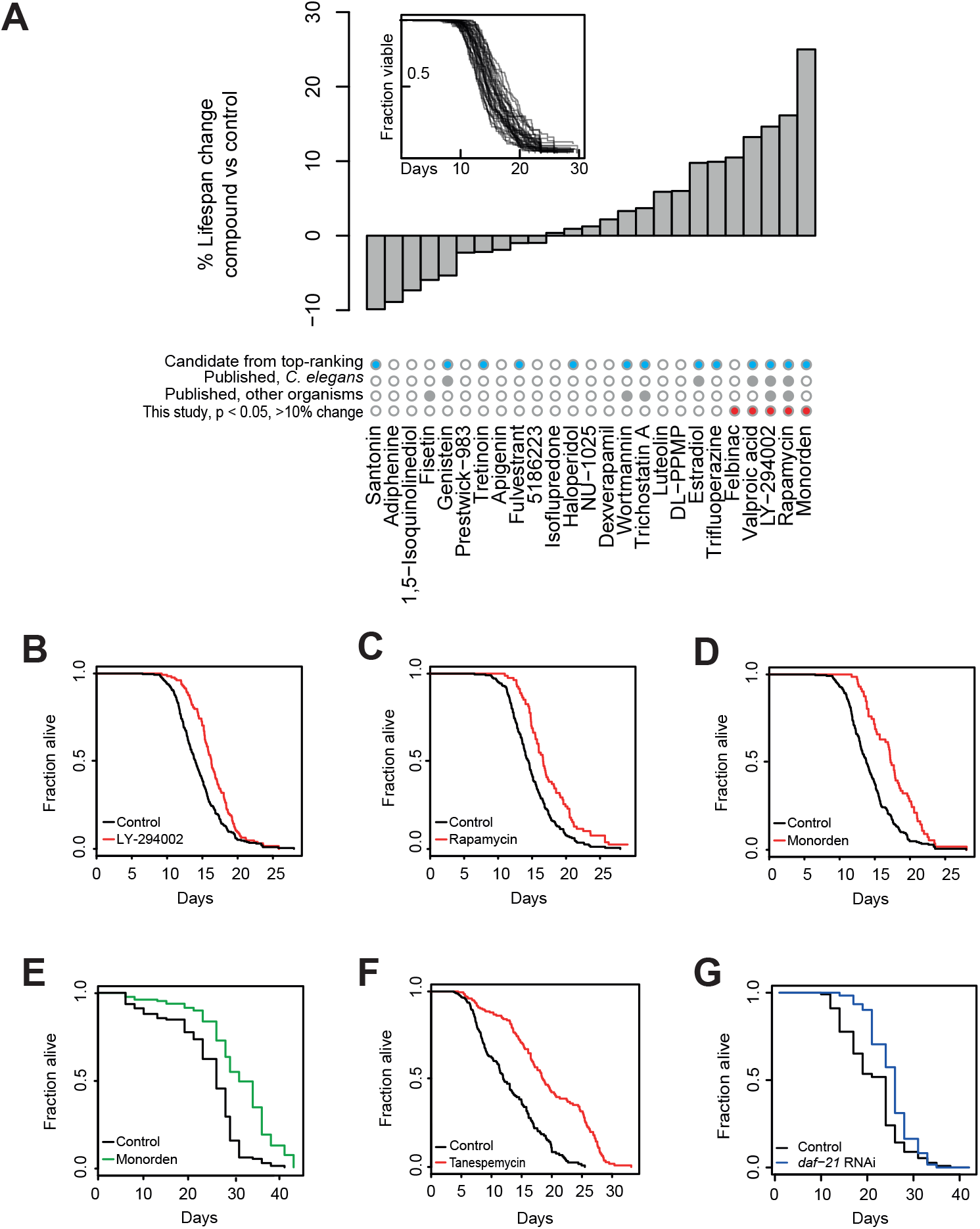
Lifespan screening and discovery of the novel geroprotective drugs Monorden and Tanespemycin. (A) High resolution lifespan curves were generated (inset) and changes in median lifespan were calculated for each candidate drug relative to the solvent control (bar graph). Compounds that were prioritized by our classification method (blue dots), that were already published to extend lifespan (grey dots), or that are significantly extending lifespan in our study (p < 0.05 and >10% extension, red dots) are indicated. (B) The survival curve of LY-294002 treated worms from (A). (C) The survival curve of Rapamycin (Sirolimus) treated worms from (A). (D) The survival curve of Monorden treated worms from (A). (E) Survival curve of Monorden treated worms, generated by using the manual scoring method of prodding with a platinum wire. (F) Survival curve of Tanespemycin treated worms. (G) Survival curve of worms grown from the L4 stage on RNAi bacteria targeting *daf-21*, the *C. elegans* gene encoding Hsp90. See Tables S8 and S10 for drug concentrations, worm numbers and statistics and Figure S3 for additional lifespan curves of compounds significantly increasing lifespan.

Notably, out of the 25 tested compounds, the mean lifespan change induced by the compounds prioritized by our classification models showed a reasonable extension (6%, Table S8), while such effect was absent from the compounds that we had selected by hand (−0.1%, Table S8). Furthermore, we observed a significant positive correlation between a better p-value and thus ranking in our geroprotector predictions and the compounds’ ability to extend lifespan in *C. elegans* (p = 0.02; Fig. S4, Table S9). And also the four compounds that exhibited the largest lifespan extension in our assays, Monorden, Rapamycin, LY-294002, and Valproic acid (Fig. 3A), were all derived from our transcriptomics-based predictions. These results underscore the power of our *in silico* approach to discover novel and, most importantly, potent geroprotective compounds.

Turning back to the actual results of our assays, we conclude that we were able to confirm the well-known geroprotective effects of Rapamycin and LY-294002, that we were able to support the lifespan extending roles of Valproic acid (which has recently been suggested to suffer from reproducibility issues (Lucanic et al., 2017)) and Tyrphostin AG-1478 (Ye et al., 2014), and that we discovered two entirely novel geroprotective compounds: Felbinac and Monorden.

### Hsp90 inhibition extends the lifespan of *C. elegans*

From here on forward, we decided to focus on Monorden (also known as Radicicol), which emerged as the most lifespan extending candidate drug from our initial assays (Fig. 3A, Table S8). Monorden is an established Hsp90 inhibitor (Griffin et al., 2004), but so far had remained unknown as a geroprotector. First, we validated its lifespan benefits by a conventional *C. elegans* lifespan assay, using manual poking (Hamilton et al., 2005). Consistent with the data from the ‘Lifespan Machine’, we again observed a robust lifespan extension by approximately 25% at 50 µM (Fig. 3E). Since Monorden is thought to target Hsp90, we next wanted to confirm that inhibition of this chaperone is indeed the mechanism by which the lifespan phenotypes are conferred. Coincidentally, we had identified the drug Tanespimycin as the third-best geroprotector candidate in our initial *in silico* screen (Fig. 2E), though had omitted it from validation in *C. elegans* due to exceptional cost of the compound. Also Tanespimycin is an Hsp90 inhibitor, though structurally quite distinct from Monorden (Blagosklonny, 2002; Schulte et al., 1998). Testing the effects of Tanespimycin at 25 μM, we observed a substantial increase in lifespan (Fig. 3F, Table S8). Further, we treated *C. elegans* with RNAi against *daf-21*, the gene encoding Hsp90 in this organism. RNAi against *daf-21* already during larval stages impairs growth and has been reported to even shorten lifespan (Somogyvári et al., 2018). Nevertheless, when we knocked down this gene only from young adulthood, animals lived longer (Fig 3G, Table S10) – which is fully consistent with our Hsp90 inhibitor data. We conclude that Hsp90 inhibition in adult *C. elegans* is a new means to extend lifespan.

### Pharmacological inhibition of Hsp90 improves the healthspan of *C. elegans*

Although living a longer life is tempting, even more importantly we would like to increase the time in our life that we spend in good health, the so-called healthspan. To test for potential healthspan benefits from Monorden treatment, we first turned again to the *C. elegans* lifespan machine (Stroustrup et al., 2013). In addition to providing lifespan data, it also generates information on the position of the worms throughout time. We reasoned that the degree of correlation of worm positions between successive timepoints provides information of the worm population’s activity level (Fig. S5). Using this type of analysis, we found that Monorden treated worms exhibited an extended period of active life, as did worms treated with the known geroprotectors Rapamycin and LY-294002 (Fig. 4A).

**Figure 4.**
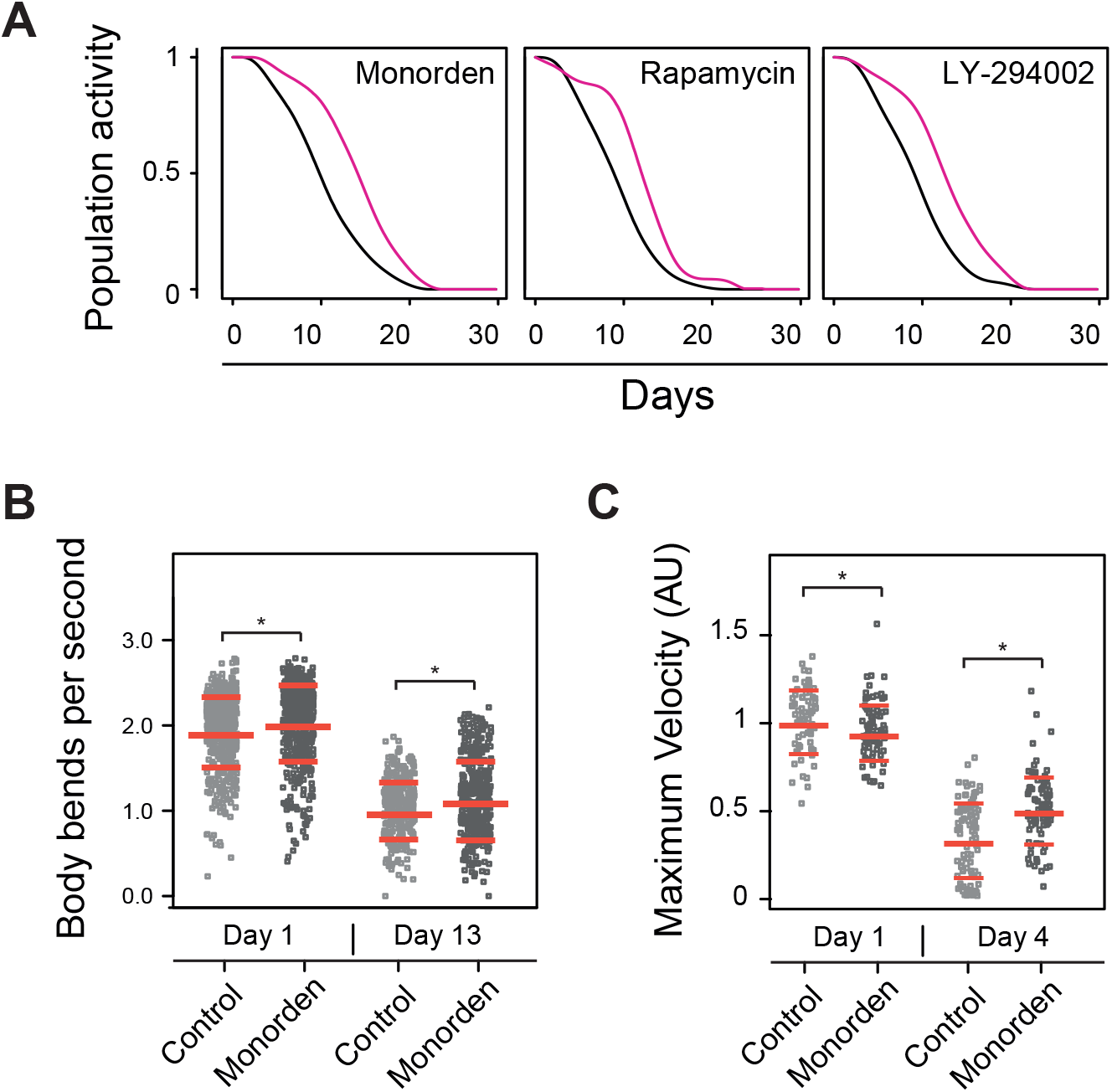
Health benefits resulting from drug treatments. (A) The activity levels of different worm populations were assessed during aging (as described in the results and further illustrated in Figure S5). The active period of life for worms treated with Monorden, Rapamycin, or LY-294002 is extended relative to controls. (B) Health as assessed by the animals’ thrashing frequency. Each dot represents a period of at least 2 seconds in a 1-minute window, during which at least 20 worms were followed. A significant increase in thrashing frequency is observed in Monorden treated worms, at both young and old ages. (C) Health as assessed by the animals’ maximum velocity in early and mid-life (day 1 and day 4 of adulthood). Each dot represents one worm’s maximum velocity during a period of 30-seconds. *: p < 0.01. See tables S11 and S12 for worm numbers and statistics.

To validate this Monorden-induced healthspan increase, we tested the motility of worms, using a ‘thrashing’ assay. This assay determines the physical motility of *C. elegans* by measuring the rate of its swimming movements in liquid. We applied Monorden to L4 worms and evaluated young (day one adult) and old (day thirteen adult) animals. Comparing young and old worms, we saw a clear decline of thrashing rate with age in control as well as treated conditions (Fig. 4B). However, Monorden treatment significantly increased thrashing rate in young and importantly also old worms by approximately 6% (p < 0.05) (Fig. 4B, Table S11).

A second metric of motility and health is the ‘maximum velocity’ of worms, being indicative of an improved healthspan especially when measured in mid-life (Hahm et al., 2015). We applied Monorden to L4 worms and measured individual worm’s maximum velocity during 30-second time frames, using young (day one adult) and mid-life (day four adult) animals. In line with our thrashing assays, we found Monorden to significantly improve animals’ maximum velocity in mid-life when compared to controls (Figure 4C, Table S12).

Taken together, these findings show that Monorden improves healthspan in addition to conferring lifespan-extending effects – a phenotype extremely desirable, if a geroprotective drug should be applied to humans for the benefit of healthy aging.

### Hsp90 inhibitors represent a novel pharmacological class of geroprotective compounds that act through the heat shock transcription factor HSF-1

Next, we wanted to gain insights into the mechanism by which Hsp90 inhibition leads to these improvements in lifespan and healthspan. We began by investigating the transcriptional response profile of Monorden and compared it to those of the established geroprotectors Rapamycin and LY-294002. From the original CMap transcriptomes, we chose three datasets generated in the same cell line (MCF7) as representative examples. Clustering analysis revealed that the effects of Rapamycin and LY-294002 are much more similar to each other than they are to the effects of Monorden (Fig. 5A, Table S13). This may be expected, since Rapamycin and LY-294002 each specifically target individual kinases in nutrient sensing pathways (Porta et al., 2014), while the effects of Hsp90 inhibitors should be much broader. In the same analysis, several transcript clusters emerged, defining the differences between the three treatments. One of these clusters stood out as being strongly upregulated upon Monorden treatment but not upon treatment with Rapamycin or LY-294002 (Fig. 5A, yellow). Using Gene Ontology (GO) term enrichment analysis, we identified this cluster to be dominated by the unfolded protein response, in particular an upregulation of heat shock proteins (HSPs) (Fig. 5B, Table S14).

**Figure 5.**
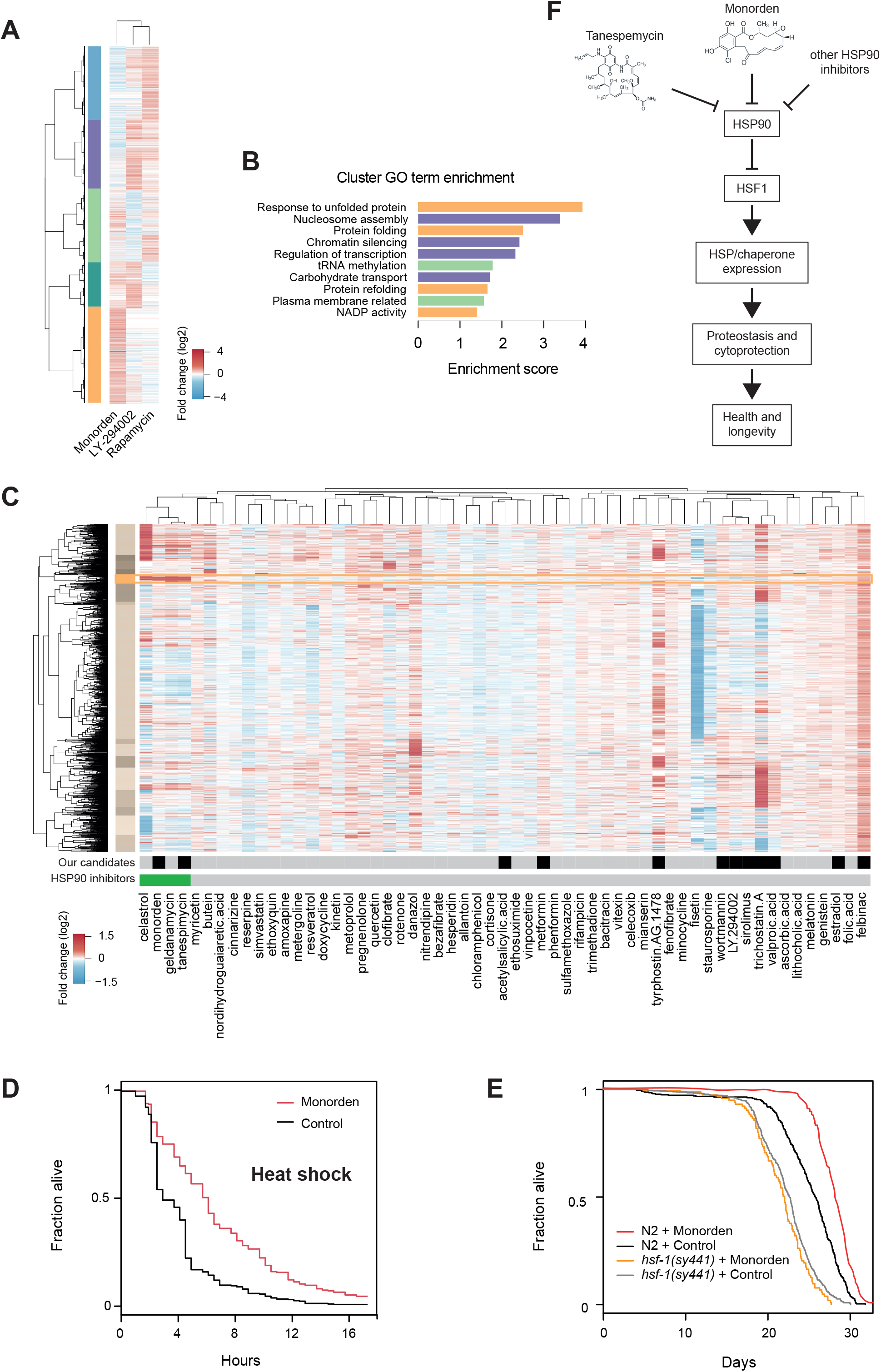
Hsp90 inhibitors and the transcriptional landscape of geroprotective drugs. (A) Heatmap clustering analysis of Monorden-, LY-294002-, and Rapamycin-induced transcriptomic changes (CMap data); only transcripts changing at least 1.5-fold are shown. Transcriptomic changes induced by LY-294002 and Rapamycin are more similar to each other than they are to the changes induced by Monorden. Five main transcript clusters emerge (blue, purple, light green, dark green, yellow). See table S13 for the genes present in the transcript clusters. (B) The ten most enriched GO terms within the five clusters from (A). They reveal that Monorden leads to a strong upregulation of the unfolded protein response, in particular an upregulation of heat shock proteins (HSPs). See table S14 for all enrichments. (C) Heatmap clustering analysis of the transcriptomic changes induced by all CMap compounds that are listed as lifespan extending in the DrugAge database, including novel compounds discovered by our study (Monorden, Tanespimycin, Felbinac). The bottom two rows indicate candidate compounds described in this study (black tiles) and any known Hsp90 inhibitors (green tiles). Only transcripts changing at least two-fold in any given drug were selected for clustering. Four compounds exhibit a substantial upregulation of HSPs (cluster highlighted by yellow box): Monorden, Tanespimycin, Geldanamycin, and Celastrol, all of which are known Hsp90 inhibitors. See Table S15 for all the transcripts used in the clustering analysis, Table S17 for the transcripts comprising the highlighted yellow cluster, and Table S16 for the GO-term enrichments within the yellow cluster. (D) Heat shock survival assay (at 35°C), showing increased viability of worms treated with Monorden relative to control. See Table S18 for worm numbers and statistics. (E) Lifespan analysis, showing that the lifespan benefits of Monorden treatment depend on the transcription factor heat shock factor 1 (HSF-1). While Monorden treatment significantly extends the lifespan of N2 animals, it fails to do so in *hsf-1(sy441)* mutant animals. See table S19 for worm numbers and statistics. (F) Model of how Hsp90 inhibitors such as Monorden and Tanespemycin may confer their geroprotective effects: 1) Hsp90 inhibitors lead to release of the transcription factor HSF1 from sequestration by Hsp90 proteins, allowing for HSF1 to trimerize and activate transcription of its target genes in the nucleus, 2) most importantly HSPs/chaperones. 3) These HSPs/chaperones help to assure the correct folding of proteins and prevent their aggregation with age, thus promoting proteostasis and cytoprotection, 4) leading to improved health and longevity.

Having established the distinguishing ability of Monorden to induce the unfolded protein response, when compared to Rapamycin and LY-294002, we then wondered how unique this ability really is, also in the broader landscape of all known geroprotectors. We thus extended our transcriptomic evaluation to all drugs that are described in DrugAge to extend lifespan and who are present in CMap. We generated transcriptomes representative of a drug response by averaging multiple instances of the same drug in CMap, and further considered only those transcripts changing by at least two-fold in any drug. We clustered these transcriptional profiles into a large heatmap, also including the novel compounds identified by our study (Fig. 5C, Table S15). To see how the unfolded protein response was modulated in this transcriptional landscape and to determine, if this was a unique ability of Monorden or Hsp90 inhibitors, we looked for transcript clusters particularly enriched for genes related to the unfolded protein response and HSP/chaperone activation. One cluster in particular caught our attention (Fig. 5C, cluster highlighted in yellow, see Table S16 for GO enrichments in this cluster). This cluster is highly enriched for genes of the unfolded protein response, including HSPs from the HSP70 class (HSPA4L, HSPA6, ASPA7) and HSP40 class (DNAJA1, DNAJC3, DNAJB6), larger (HSPH1) and small (HSPB1) HSPs, among others (Table S17). Figure 5C revealed that only four compounds are clear activators of this transcript cluster. Strikingly, all of these are known Hsp90 inhibitors: Monorden, Tanespimycin, Celastrol, and Geldenamycin. Celastrol is a pentacyclic triterpenoid (Hieronymus et al., 2006), described to induce heat shock response (Trott et al., 2007; Westerheide et al., 2004) and previously shown to increase the lifespan of *C. elegans* by 17% (Jung et al., 2014). Geldanamycin is a 1,4-benzoquinone ansamycin antibiotic. In *C. elegans*, a prior studies failed to observe robust lifespan and healthspan benefits from this drug (Calvert et al., 2016). However, it has been argued that Geldanamycin actually fails to bind specifically the *C. elegans* Hsp90, due to it harboring minor structural differences from the human protein (David et al., 2003). In the light of the mostly side-effect-free lifespan benefits of the other three Hsp90 inhibitors, we thus suggest that Geldanamycin might indeed not function as an Hsp90 inhibitior in *C. elegans* and instead causes off-target effects. Taken together, the above observations show that Hsp90 inhibitors generally lead to an upregulation of HSPs/chaperones, a feature that is unique and defines them as a new pharmacological class amongst geroprotectors.

It is well established that increased stress resistance can slow down the aging process and thereby improve organismal health and longevity. HSP upregulation confers such stress resistance by improving the organism’s protein homeostasis. We therefore hypothesized that HSP upregulation comprises the mechanism by which Hsp90 inhibitors confer their beneficial effects on health and lifespan. To confirm this, we first tested whether a process that is particularly dependent on HSPs, namely the resistance to heat-induced unfolded protein stress, can be improved by Monorden. Thus, we grew *C. elegans* at 20°C, exposed them to Monorden from the L4 stage onwards, and on day one of adulthood shifted them to 35°C. In line with our hypothesis, we found that Monorden-treated animals survived significantly longer under these conditions (Fig. 5E, Table S18). This observation shows that Monorden not only extends lifespan but also improves resistance to heat stress, an ability that is well known to depend on HSP induction (Wu, 1995). Further, it illustrates that upregulation of HSPs by Monorden is functionally relevant and may indeed be the key mechanism of its geroprotective effects. To further support this notion, we turned to evaluation of the conserved transcription factor HSF-1. HSF-1 is a master regulator of HSP expression in eukaryotes, required for the upregulation of HSPs upon unfolded protein stress. Overexpression of HSF-1 and the resulting HSP induction have been found sufficient to increase lifespan in worms (Hsu et al., 2003). We therefore tested, whether a hypomorphic allele of HSF-1, *hsf-1(sy441)*, would be able to suppress the lifespan extension phenotype caused by Monorden. Indeed, this allele led to a full suppression of the lifespan benefits caused by Monorden (Fig. 5F, Table S19). Since it has been well established that Hsp90 functions as an inhibitor of HSF-1 (Zou et al., 1998), our results eventually led us to propose the following mechanistic cascade that would lead to prolonged lifespan and health in *C. elegans*: Compounds like Monorden or Tanespemycin cause inhibition of Hsp90, which in turn causes activation of HSF-1. HSF-1 as a transcription factor then upregulates HSP expression, leading to improved protein homeostasis throughout age and thereby the compounds’ geroprotective effects (see also Fig. 5F). Notably, this mechanism also demonstrates for the first time that aging-preventive effects of HSF-1 can actually be activated by pharmacological means, namely by targeting its upstream regulator Hsp90.

### Hsp90 inhibition shows good utility as a combinatorial treatment with other geroprotectors

By now we had established that Hsp90 inhibitors lead to rather distinct transcriptional responses including a unique ability to upregulate HSPs, when compared to other geroprotectors in our transcriptome clustering analysis (Fig. 5C). Thus, we hypothesized that Hsp90 inhibitors may provide excellent complementing capabilities, when used in combinatorial treatments with other geroprotectors, resulting in additive beneficial effects on lifespan. To test this, we treated worms with 50 μM of Monorden, Rapamycin, or LY-294002 as well as combinations of 50 µM Rapamycin + 50 µM Monorden or 50 µM LY-294002 + 50 µM Monorden (Fig. 6, Table S20). Indeed, Monorden treatment was able to significantly extend the lifespan of either Rapamycin or LY-294002 treated animals – by an additional ~20%. In contrast, increasing the dose of Monorden alone did not lead to an additional lifespan extension (Fig. S6, Table S20). In summary, Monorden appears to target geroprotective pathways that are at least in part distinct from those targeted by Rapamycin and LY-294002, which highlights Hsp90 inhibitors not only as good geroprotectors when used by themselves, but also compounds that may further enhance the beneficial effects of other geroprotectors by targeting distinct but complementing pathways.

**Figure 6.**
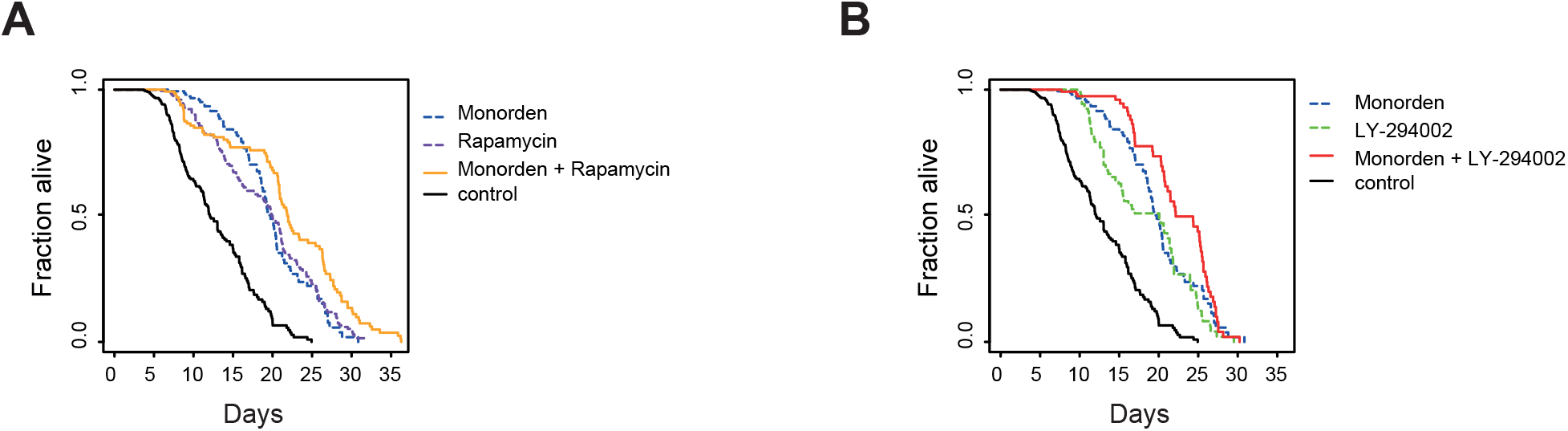
Additive effects of Monorden treatment with other geroprotective compounds. (A) Lifespan assay of worms treated with either solvent control, Monorden alone, Rapamycin alone, or the combination of Monorden and Rapamycin. The latter combinatorial treatment shows an increased lifespan benefit (p < 0.01), suggesting that Monorden and Rapamycin target different longevity pathways. (B) Lifespan assay of worms treated with solvent control, Monorden alone, LY-294002 alone, or the combination of Monorden and LY-294002, likewise shows additive benefits of the compounds (p < 0.01). See table S20 for worm numbers and statistics.

## Discussion

We have established here a novel and powerful strategy for the discovery of geroprotective compounds, using age-classifiers derived from age-stratified human tissue transcriptomes and applying these to a database of drug-induced transcriptomic changes in human cell cultures. We reasoned that our strategy would be able to identify drugs that produce a ‘youthful’ transcriptional signature and thus reveal candidate geroprotective compounds. Indeed, when we applied this strategy to datasets from the GTEx consortium and CMap and generated short-lists of geroprotector candidates, we were able to validate many of our top hits as being lifespan-extending in the model organism *C. elegans*. Specifically, we confirmed the lifespan-extending abilities of known geroprotectors like Tyrphostin AG-1478, LY-294002, and Rapamycin. More importantly however, we identified novel geroprotectors that never before had been described to prevent aging or extend the lifespan in any organism, namely Felbinac, Monorden, and Tanespimycin. By eventually focusing on both, Monorden and Tanespimycin, we reveal the existence of a new and very potent class of geroprotectors that act via Hsp90 inhibition and concomitant induction of HSP expression.

Hsp90 is a chaperone protein, comprised of an N-terminal nucleotide binding and ATPase domain, a middle domain involved in the binding of its substrates, and a C-terminal domain involved in Hsp90 dimerization and its association with co-chaperones (Powers and Workman, 2007; Welch and Feramisco, 1982; Welch et al., 1991). It accounts for about 1-2% of all proteins in the cell (Powers and Workman, 2007; Welch and Feramisco, 1982; Welch et al., 1991) and together with co-chaperones and Hsp70 forms a sophisticated chaperone machinery (Pratt and Toft, 2003). Specifically, Hsp90 serves to fold, regulate, or sometimes also to aid in the degradation of over 200 so-called ‘client proteins’. These client proteins are enriched for both protein kinases and transcription factors (Pratt and Toft, 2003; Trepel et al., 2010). Several of these have been found to be growth promoting or even oncogenic, including AKT, mTOR, ERBB2, BCR-ABL, C-RAF, CDK4, various steroid hormone receptors, and telomerase (Blagosklonny, 2002; Echeverría et al., 2011; Powers and Workman, 2007). As a consequence, Hsp90 has been widely acknowledged as a therapeutic target for cancer (Calderwood et al., 2006; Whitesell and Lindquist, 2005) and has been intensely subjected to drug development efforts (Powers and Workman, 2007). By now, pharma companies have filed over 30 patents for compounds that target Hsp90 (Sidera and Patsavoudi, 2014). Pharmacological impairment of Hsp90 can occur in a variety of ways: Most Hsp90 inhibitors block the ATP binding site of Hsp90 (e.g. Monorden, Tanespimycin, and Geldanamycin) (Prodromou et al., 1997; Schulte et al., 1998; Taipale et al., 2010), but others can also disrupt co-chaperone or client interactions (e.g. Celastrol (Chadli et al., 2010)), or interfere with post-translational modifications of Hsp90 (Li et al., 2009).

The first described inhibitors of Hsp90 were naturally occurring compounds, including Geldanamycin and Monorden. Clinical exploration of Geldanamycin was halted however, due to toxic side effects. This is also consistent with a recent study in which this Hsp90 inhibitor showed negative side-effects in *C. elegans* (Calvert et al., 2016). Eventually, structural analogs of Geldanamycin with lower toxicity were developed (Blagosklonny, 2002; Supko et al., 1995). Tanespimycin is such an analog, which has already led to more promising results in clinical trials (Banerji, 2009; Pacey et al., 2006; Sharp and Workman, 2006). Structural variants of Monorden that improve its potency and stability have also been developed with promising results (Soga et al., 1999), and our work prompts their consideration as geroprotectors in the future.

Besides disrupting growth-promoting protein kinase signaling and other oncogenic pathways, Hsp90 inhibitors have a well-documented ability to induce the expression of heat shock proteins (HSPs) (Clarke et al., 2000; McCollum et al., 2006). This induction is thought to occur through the heat shock transcription factor HSF1, a client protein of Hsp90 (Trepel et al., 2010). Upon Hsp90 inhibition, HSF1 becomes activated by dissociating from Hsp90 and undergoing trimerization as well as nuclear translocation. Once in the nucleus, HSF1 promotes the expression of heat shock response genes including HSPs/chaperones (Zou et al., 1998). In the context of cancer therapy, this is an undesirable side effect of Hsp90 inhibitors, since HSPs may serve to protect cancerous cells. To avoid these consequences, HSF1 repressors have been developed for combinatorial use with Hsp90 inhibitors (Trepel et al., 2010). In the context of aging however, this activation of HSF1 should be beneficial. Consistently, overexpression of HSF1 has been shown to increase lifespan in *C. elegans* (Baird et al., 2014; Hsu et al., 2003), and we show in Figure 5E that the lifespan-extending benefits of Hsp90 inhibition are abrogated by removal of HSF1 function. Taken together, we have identified Hsp90 inhibition as a potent strategy to improve healthspan and lifespan, through the transcription factor HSF1 and presumably its ability to induce the expression of HSPs.

Interestingly, while Hsp90 inhibition leads via this mechanism to already remarkable geroprotection in *C. elegans*, there are reasons to believe that the geroprotective benefits in humans should even be greater. Previous work has shown that there are two hallmarks of human aging, which don’t exist in *C. elegans*, that additionally can be targeted by Hsp90 inhibition: 1) chronic inflammation and 2) the presence of senescent cells. Hsp90 inhibitors are able to suppress inflammation by blocking immune cell activation (Tukaj and Węgrzyn, 2016), and a recent screen identified Hsp90 inhibitors to be senolytic agents (Fuhrmann-Stroissnigg et al., 2017). Thus, it can be assumed that Hsp90 inhibition confers geroprotection through even multiple mechanisms in humans, involving improved protein homeostasis (Conte et al., 2008; Griffin et al., 2004; Sonoda et al., 2010), as described in our study, but likely also through decreased inflammation (Zhao et al., 2013), and the removal of senescent cells (Fuhrmann-Stroissnigg et al., 2017).

Finally, our work supports the idea that Hsp90 inhibitors may be particularly useful in combinatorial treatments with other geroprotectors. By transcriptome analyses from human cell culture we have demonstrated the distinct and thus complementary nature of the gene expression changes caused by Hsp90 inhibitors, when compared to other known geroprotective drugs, and by using *C. elegans* we have shown *in vivo* the additive longevity effects that result from Hsp90 inhibition when combined with mTOR or PI3Kinase inhibition. These observations encourage the further evaluation of Hsp90 inhibitors in combination with other geroprotective compounds for additive beneficial effects in humans.

In summary, our study provides a new state-of-the-art approach to discovering geroprotective compounds, and highlights Hsp90 inhibition as a promising new therapeutic approach to defer aging and age-related complications. Several well-developed Hsp90 inhibitors are already available that could be repurposed for such use, and future work in humans will have to determine their full potential.

## Experimental Procedures

### Pre-processing of the GTEx data

The publicly available Version 6 of the GTEx Transcriptome datasets (accessed April 2016, dbGaP Accession phs000424.v6.p1) and annotation files (subject and sample attributes) were downloaded from the GTEx portal (https://gtexportal.org/home/datasets). Transcriptome data was loaded using the ‘read.gct’ function in the ‘CePa’ package (Centrality-based pathway enrichment) (Gu and Wang, 2013) into R. Data was preprocessed as previously described (Mele et al., 2015; Taskesen and Reinders, 2016; The GTEx Consortium, 2015): Data was normalized to 1 million reads, low abundant transcripts were removed by selecting RPKMs whose values were less than 0.1 in at least 80% of samples. A pseudocount of 1 was added and data was log2-transformed. Transcriptomes were filtered to contain only those with an RNA integrity score (RIN) greater or equal to 6 (the SMRIN column of the annotation file) and for those that were deemed usable by the GTEx consortium (the SMAFRZE column of the annotation file). For data processing, GTEx files were parsed into sub files by tissue. In summary this resulted in a dataset of 8555 transcriptomes covering 19343 transcripts in 51 tissues, both genders, and grouped into decade-sized age bins.

### Processing of the GTEx data for age-related comparisons

Genders were treated separately, and all tissues were systematically processed in the same way. Tissues were individually loaded and parsed into age bins. Old was considered to be the age bin of 60-69 (the age bin of 70-79 was omitted due to low sample numbers), and young was considered to be any of the age bins of 20-29, 30-39, 40-49, or 50-59, which then would be compared to the old age-bin in a binary fashion. Young age groups were compared to the old, only when both groups contained at least 10 transcriptomes each. Next, the following steps were performed for the binary comparisons: Transcriptomes were reduced to only contain transcripts in common between the GTEx and CMap datasets (10,890 transcripts). Transcriptomes were further filtered to remove low abundant transcripts (the 10% least abundant). To aid the model building, a feature reduction step was undertaken by keeping only those transcripts which showed differential expression between the young and old age bin (p < 0.01). This was achieved by the ‘rowttests’ function in the R package ‘genefilter’ (Gentleman et al., 2017). Any cell line-based data in GTEx was omitted from our study.

### Random Forest model generation and selection

For each binary comparison of ‘old’ transcriptomes to any given ‘young’ transcriptomes prepared above, data was reduced to have equal numbers of samples in each age group. Data was split into 70% used for model generation and 30% reserved for model validation, utilizing the ‘createDataPartition’ function in the R package ‘caret’ (Kuhn et al., 2012). The 70% data partition was centered and scaled using the ‘preProcess’ function of the same R package. Next, the R package ‘Random Forest’ (Liaw and Wiener, 2002) was implemented using the ‘train’ function again in the ‘caret’ package and was used to train models, using repeated cross validation (10 fold), using the default setting of 500 trees per forest, and using a standardized grid that varied the number of variables randomly selected per data split (varying .mtry in the tunegrid parameter). Models were automatically tuned, selecting the parameters with the best performing receiver-operating characteristic (ROC). Final models were tested on the remaining 30% data split for accuracy. 182 models were generated in this way. Performance of the models on the testing dataset was assessed using the ‘performance’ function in the R package ‘ROCR’ (Sing et al., 2007). Final models were evaluated based on the ROC, the sensitivity, and the specificity achieved during model training, and the ROC and accuracy determined during model testing. Only those passing cutoff criteria above 0.75 in all these domains were retained. This resulted in 24 final models, our age-classifiers, able to distinguish young transcriptomes from those in the old age group of 60-69 years. Fourteen models derived from male tissues, from the adrenal gland (age 40-49), the coronary artery (age 40-49), mammary breast tissue (age 40-49), the transverse colon (age 40-49), the gastroesophageal junction of the esophagus (age 40-49), the pancreas (age 40-49 and also 30-39), the aorta artery (30-39), the esophagus muscularis (age 30-39), suprapubic skin, not sun exposed (age 30-39 and also 20-29), the thyroid (age 30-39), and the prostate (age 20-29). Ten models derived from female tissues, from the adrenal gland (age 50-59, and also 40-49), the coronary artery (age 50-59, and also 40-49), the liver (age 50-59), the pituitary (age 50-59), the vagina (age 50-59, and also 40-49), the gastroesophageal junction of the esophagus (age 40-49), and the heart’s atrial appendage (age 40-49).

### Connectivity Map data preparation

Information on the Connectivity Map (CMap) instances (drug perturbation IDs and descriptions) was accessed through the R package ‘ConnectivityMap’ (Package and Shannon, 2013). A corresponding matrix of ‘amplitudes’ (a measure of the extent of differential expression of a given probe set) was obtained by download from the Broad Institute’s FTP server (ftp://ftp.broadinstitute.org/distribution/cmap). The CMap amplitude matrix was converted to a fold change (log2) matrix using the conversion equation provided at http://www.broadinstitute.org/cmap/help_topics_frames.jsp. In order to later apply the CMap fold change data to the GTEx dataset, we converted the affymetrix probe IDs used in CMap to Ensemble gene IDs (used in GTEx) through the R package ‘biomaRt’ (Durinck et al., 2005, 2009). For each GTEx gene entry, the corresponding probe fold change entries from CMap were collapsed to one value (if needed) in a conservative fashion by taking the median entry. Of the 19434 transcript entries in GTEx, we found 10869 to have corresponding CMap entries. Finally, the data was converted to linear scale, which resulted in a CMap dataset of fold changes that could be applied to transcriptomes from the GTEx dataset.

### Generating ‘drug-induced’ transcriptomes for age classification

For each specific model, derived from a gender, tissue, and binary age group comparison within the GTEx dataset, the transcriptome of a corresponding ‘prototypical’ age was generated, representing an average transcriptome between the compared old and young datasets. This was done by first determining the median transcriptional profile in the young and old datasets separately and subsequently averaging the two. This resulted in 24 prototypical transcriptomes, one for each model, which we termed ‘middle aged’. The processed CMap fold changes were then applied to each of these middle-aged transcriptomes, generating datasets that we term ‘drug-induced’ transcriptomes, which are specific for each model. These transcriptomes are then ready to be used for age classification.

### Application of age-classifiers to ‘drug-induced’ transcriptomes and the eventual ranking of geroprotectors

For each model, the corresponding ‘drug-induced’ transcriptome dataset was preprocessed by scaling and normalizing with the relevant model’s parameters, and were classified by the respective models, returning a probability score of being ‘young’. This resulted in the 6100 CMap perturbation transcriptomes receiving a classification from each of the 24 models. For each drug present in replicate or various conditions in CMap, only single score was used per model, namely the score from the most geroprotective prediction. This resulted in each drug being assessed for its geroprotective potential 24 times (once for each model), regardless of the number of replicates or conditions in CMap. To compare between models, these predictions were normalized to a maximal value of 0.5 and subsequently centered around 0. A drug was given a final geroprotective ranked score, by seeing how many of the 24 predictions placed the drug above the mean absolute deviation of the overall distribution of all drug scores, and generating a p value using the hypergeometric function, with a Benjamini correction.

### DrugAge

The DrugAge database (Barardo et al., 2017) was accessed in May 2017, and was filtered to include all drugs that significantly increase lifespan.

### Gene ID conversion

Where needed and unless otherwise specified, to convert between Ensemble Stable Gene IDs and common names, the BIOMART website (http://www.ensembl.org/biomart) was used, with the Human genes dataset (GRCh38.p10) for conversion.

### GO term enrichments

Gene functional enrichments were determined by using the DAVID Bioinformatics Resources (version 6.8) (Huang et al., 2009). Corresponding background gene lists of indicated size (see source datasets provided) were used for each enrichment analysis. Annotation clusters determined by DAVID (groupings of related genes based on the agreement of sharing similar annotation terms) having an enrichment score of >1 were selected for consideration. A representative naming for the enrichment was selected after evaluation of the annotation cluster’s GO terms.

### Data Visualizations

Data was visualized using R (R Core Team, 2013) with color schemes selected from there or the RColorBrewer package (Neuwirth, 2014).

### Worm strains and maintenance

*Caenorhabditis elegans* were maintained on NGM plates (Stiernagle, 2006) and fed with *Escherichia coli* OP50-1 unless stated otherwise. *C. elegans* strains used in this study were N2 (wild type) and PS3551 (*hsf-1(sy441)*).

### RNAi by feeding

*C. elegans* were grown on the *E. coli* strain HT115 without plasmid until the L4 stage, then washed in M9 buffer containing antibiotics (Tetracyclin, Carbenicillin, and Streptomycin) to remove these bacteria, and finally transferred to HT115 containing dsRNA-expressing plasmids targeting *daf-21*. These bacteria were obtained from published collections (Kamath et al., 2003; Reboul et al., 2003). HT115 containing empty plasmid L4440 was used as a negative control.

### Compounds

The following compounds were purchased from Sigma Aldrich: Dimethyl sulphoxide (DMSO, ref. D2650), Valproic acid (ref. S0930000), Trichostatin A (ref. T8552), Tanespimycin (ref. A8476), Fulvestrant (ref. I4409), LY-294002 (ref. L9908), Estradiol (ref. E8875), Rapamycin (ref. 37094), Haloperidol (ref. H1512), Prochlorperazine (ref. P9178), Genistein (ref.G6649), Trifluoperazine (ref. T8516), Santonin (ref. Y0001052), Tretinoin (ref. R2625), Monorden (ref. R2146), Wortmannin (ref. W1628), 1,5-isoquinolinediol (ref. I138), Dexverapamil (ref. V4629), Fisetin (ref. F4043), Luteolin (ref. L9283), Cantharidin (ref. C7632), DL-PPMP (ref. P4194), Tyrphostin AG-1478 (ref. T4182), Apigenin (ref. A3145), Adiphenine (ref. A3649), Isoflupredone (ref. 1348907), Felbinac (ref. Y0000731), Prestwick-983 (ref. D7571), NU-1025 (ref. N7287). The compound ‘5186223’ was purchased from ChemBridge. Compounds were dissolved in water, ethanol, or DMSO, in concentrations as specified (Table S8). For all compounds dissolved in DMSO, DMSO concentration in drug and control plates was kept at 0.33% (v/v), as recommended by several studies (Calvert et al., 2016; Ye et al., 2014). Ethanol concentrations were likewise kept at 0.33% (v/v). Two compounds, Fulvestrant and 5186223 required higher final DMSO concentrations (0.66% (v/v)) due to solubility issues.

### Preparation of drugged NGM plates

NGM plates for lifespan and healthspan assays were identical to the maintenance plates mentioned above, with two exceptions: 1) Ampicillin (100 μg/ml) was added to the plates, and 2) the plates were seeded with dead instead of live OP50-1 bacteria, to avoid secondary effects from bacterial drug metabolism. The bacteria were killed by heat. Finally, drugs or solvent controls were added to the plates, the plates then incubated over night at room temperature, before the worms were added.

### Lifespan assays

Synchronized worms were grown on standard NGM plates and fed with live OP50-1 at 20°C until the late L4 stage, washed, and then transferred to drugged or solvent control NGM plates. Worms were divided among at least 6 plates (3 cm) per condition, with 10-25 worms per plate, and plates were treated with 0.1 g/ml 5-Fluoro-2′-deoxyuridine (FUDR) to prevent progeny production (Hamilton et al., 2005). Their survival was recorded and analyzed by a fully automated ‘Lifespan Machine’, as previously described (Stroustrup et al., 2013). To ensure high quality results, lifespan curves were assessed, only if they were derived from least 50 tracked worms. Data assessment was conducted using the ‘Survival’ package in R (Therneau, 2012) with Kaplan-Meier estimated survival curves compared to each other using the log-rank test. For the lifespan experiment shown in Fig. 3E and G, identical procedures were followed, except that survival of animals was examined manually every 2-3 days as previously described (Hamilton et al., 2005).

### Heat-stress assay

Synchronized N2 animals were grown and transferred to drugged or solvent control NGM plates as described for lifespan assays. At day 1 of adulthood, animals were shifted to 35°C, and their survival was recorded and analyzed by the ‘Lifespan Machine’ as previously described (Stroustrup et al., 2013).

### Healthspan analysis: Population activity assay

To derive ‘activity curves’ of worm populations as a healthspan measure from our lifespan machine data, we utilized the ‘animal_position.csv’ output file that contains positions of ‘worm objects’ detected by the software. These are objects that the software presumes to be worms. We reconstructed the image space by creating a binary matrix corresponding to ‘non-worm’ (0) and ‘worm’ (1) positions, as indicated by the file, and as plotted in Figure S5A. Converting this binary image matrix to a linear vector allowed for Pearson’s correlations between time points to be calculated. We compared in this way every time point (n) with the one occurring directly subsequent to it (n+1) (see Fig. S5A for illustration), generating a time-based correlation of worm positions (Fig. S5B). A higher correlation between two adjacent time points was interpreted as less movement of the worms, indicative of lower ‘health’ of the population. Data from timepoints beyond the death of the worm population were omitted, to avoid artifacts. Combining these correlations from independent NGM plates, generated a large representative plot of time-based correlations, reflecting the worm population’s activity in time on the plates (Fig. S5C). Fitting these correlations with a linear smoothing spline, using the preinstalled smooth.spline function in R, produced a curve that could be interpreted as movement of worms on the plates at the population level (Fig. S5D). We subsequently made simple assumptions to plot this as a more familiar ‘curve’ for visual comparison between drug treatments and controls. Namely, 1) we mirrored and normalized the correlations, scaling them from 1 (‘most active’) to 0 (‘least active’), where we assumed that the highest correlation was the ‘least active’ time point and the lowest correlation was the ‘most active’ time point. 2) We assumed that the ‘most active’ time point represented the population optimum, and set all previous values to this. Further, we assumed that the ‘least active’ time point represented the end of movement on the plate, and set all subsequent time points to that value. Performing this for drug treatment and control conditions, allowed to assess a compound’s influence on the activity of a worm population as a measure of the population’s healthspan (Fig. 4A).

### Healthspan analysis: Maximum velocity assay

Individual worms’ maximum velocity was measured as previously described (Hahm et al., 2015). In brief, synchronized N2 animals were grown and transferred to drugged or solvent control NGM plates, as was done for lifespan assays. On day 1 and day 4 of adulthood, individual worms were transferred to individual NGM plates without bacteria. The worms’ movement was video-recorded immediately for 30 seconds (10 frames per second), and the velocity profile was analyzed using the ImageJ plugin ‘wrMTrck’ (Nussbaum-Krammer et al., 2015) and Microsoft Excel. Population distributions were assessed using the Wilcoxon-Mann-Whitney test from the preinstalled ‘wilcox.test’ function in R.

### Healthspan analysis: Thrashing assay

Synchronized N2 animals were grown and transferred to drugged or solvent control NGM plates, as was done for lifespan assays. As the only difference, here 24-well plates were used. On day 1 and day 13 of adulthood, worms were measured for body bend movement in liquid. In brief, M9 was added to individual plate wells and videos were immediately recorded for a duration of 2 minutes (at 20 frames per second). Body bends were measured using the ImageJ plugin ‘wrMTrck’ (Nussbaum-Krammer et al., 2015). Population distributions were assessed using the Wilcoxon-Mann-Whitney test from the preinstalled ‘wilcox.test’ function in R.

## Author contributions

G.E.J., X.X.L., E.A.A.N., and C.G.R. conceived and designed the analyses and experiments. G.E.J. conducted the bioinformatic analyses. X.X.L., G.E.J., R.I.S., and L.M.A. conducted the *in vivo* experiments and analyzed the resulting data. N.S. helped with the setup of the ‘lifespan machine’. G.E.J., X.X.L., and C.G.R. wrote the manuscript.

## Acknowledgements

We thank Peter Swoboda for advice and infrastructural support as well as João Pedro de Magalhães for providing early access to the DrugAge database. We thank Maria Eriksson and Urban Lendahl for comments on the manuscript. C.G.R was supported by the VR grant 2015-03740, the GENiE COST grant BM1408, and an ICMC project grant.

## Supplementary Figures

**Figure S1.**
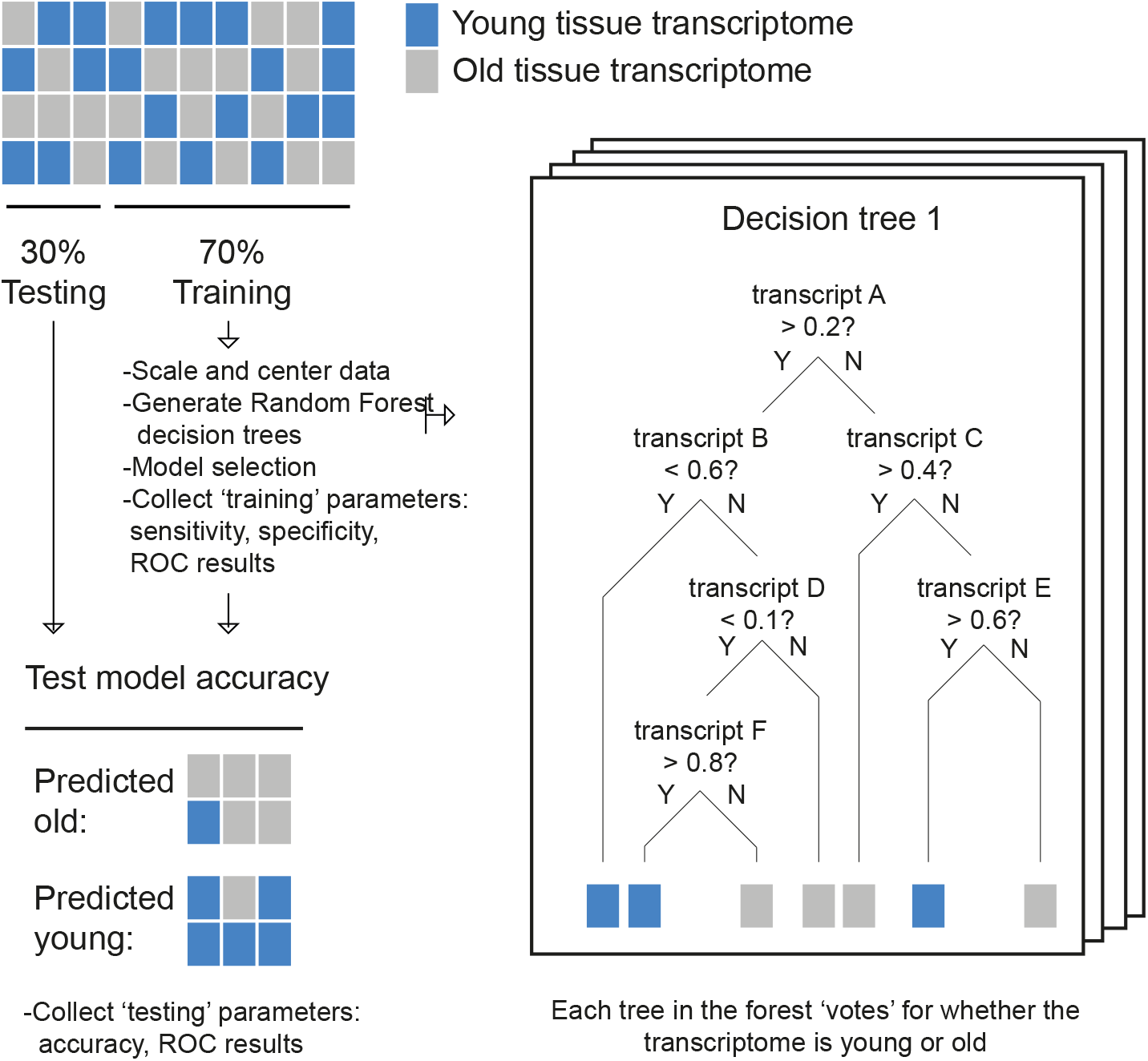
Generation of Age Classifying Models. Prior to age classification, the GTEx transcriptome datasets are separated by tissue, gender, and age into decade-wide age-bins, requiring at least 10 samples per bin. A particular ‘young’ and ‘old’ bin pair is selected, where the tissue and gender are identical, and where ‘old’ is defined as the 60-69 age bin, and ‘young’ is any of the younger age bins. Pairwise comparisons of transcripts between ‘young’ and ‘old’ are performed only on differentially expressed transcripts (p<0.01) to reduce the dataset and aid the classification. Data is subsequently down sampled to have equal numbers of old and young samples per comparison. 70% of the GTEx data is used to train models, by scaling and centering the data and eventually growing 500 random forest decision trees. Parameters of sensitivity, specificity, and ROC (receiver operating characteristic) are recorded in this ‘training’ phase. The models are tested against the remaining 30% of the GTEx data in the ‘testing’ phase to determine their accuracy and ROC in the independent data partition. Final models are selected, based on results from both, the training and testing phases.

**Figure S2.**
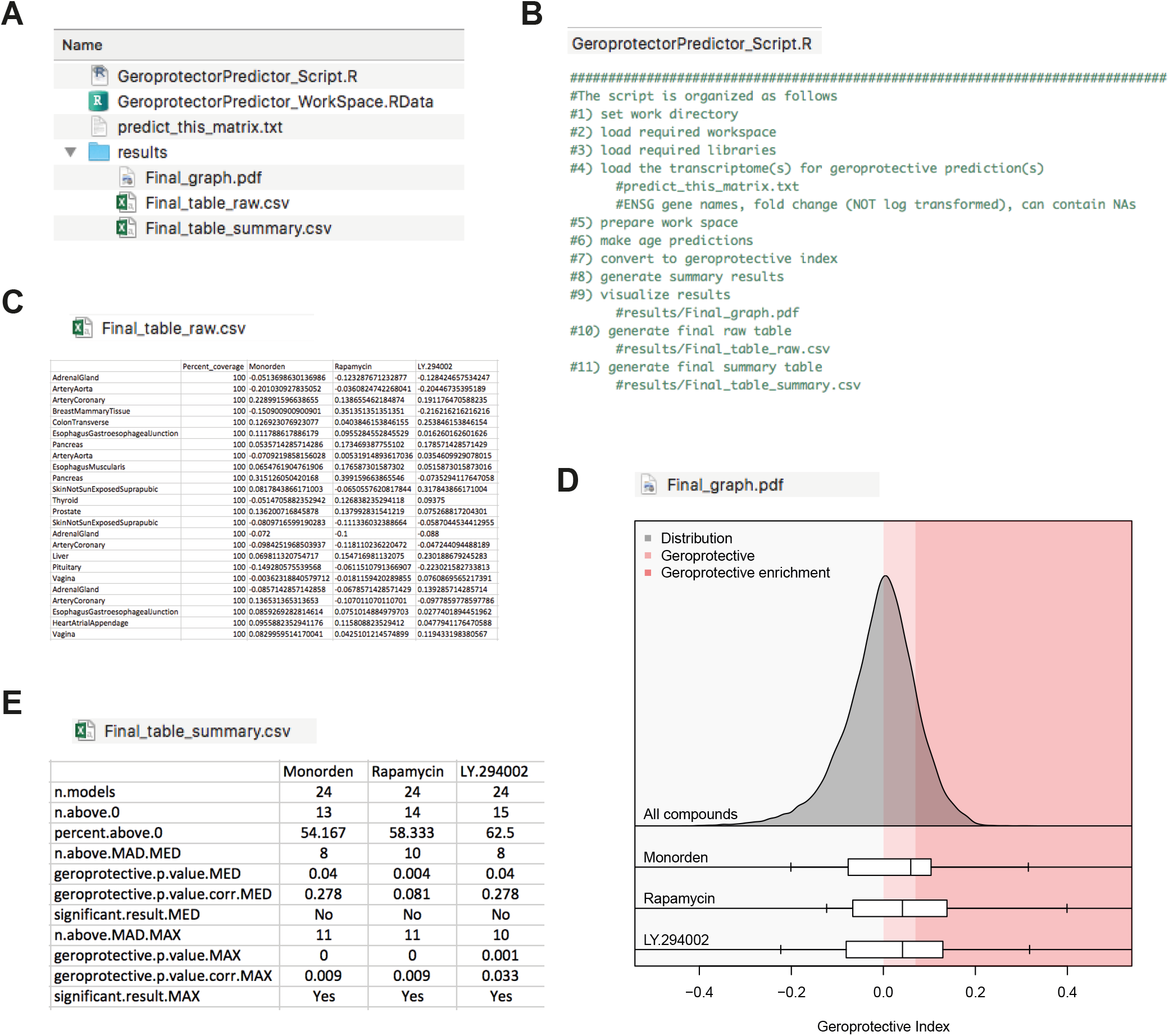
Pipeline for the transcriptomics-based identification of geroprotector candidates. Accompanying this manuscript, a ‘stand-alone’ script is provided, which can be used to determine the geroprotective index scores of unknown compounds, based on their transcriptomic effects. The script is provided as a compressed file (GeroprotectorPredictor.zip). Unpackaging it provides (A) an R script, all required data (models, variables, etc), and a modifiable txt file that can be used to input transcriptome(s). (B) An outline of the steps in the R script is provided. Running the R script produces results that are stored in a results folder. These include (C) the raw predictions from each model as geroprotective index scores, (D) a final graph showing the predictions of the input transcriptomes relative to all other predictions from CMap, and (E) a summary table providing final results, most importantly enrichment scores and their significance.

**Figure S3.**
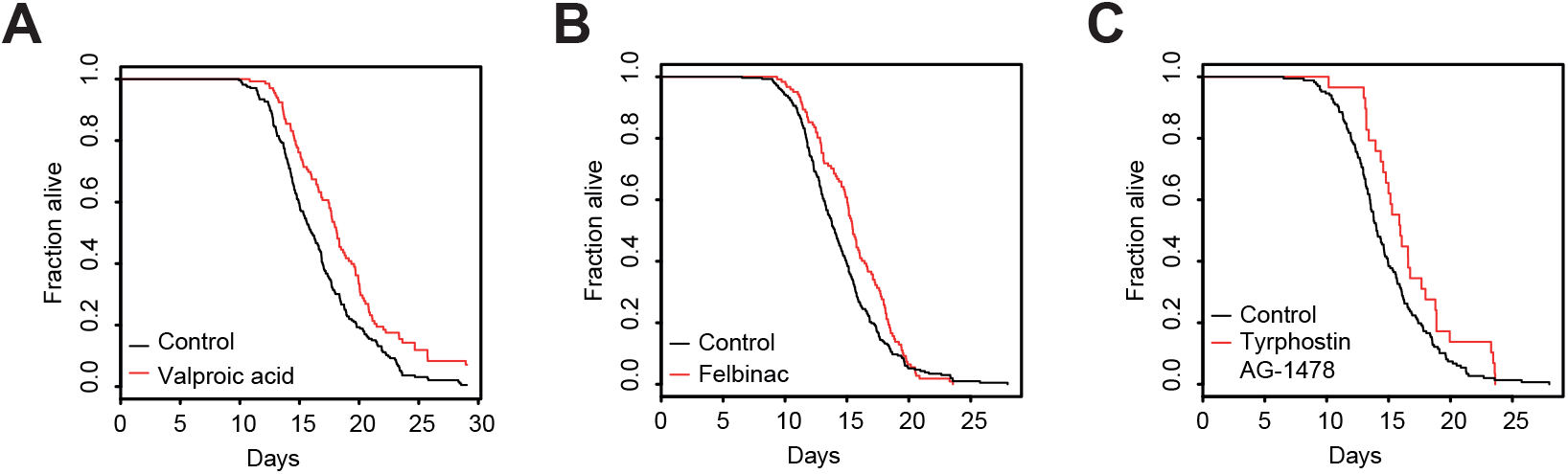
Lifespan curves of additional candidates. Survival curves of additional candidate compounds, showing a greater than 10% lifespan extension with a significance of p < 0.05. (A) The survival curve of Valproic acid treated worms. (B) The survival curve of Felbinac treated worms. (C) The survival curve of Tyrphostin AG-1478 treated worms. See Table S8 for drug concentrations, worm numbers and statistics. Tyrphostin AG-1478 results were omitted from Figure 3A due to insufficient worms in the first lifespan experiment (<50), though results were later confirmed with higher worm numbers (see Table S8).

**Figure S4.**
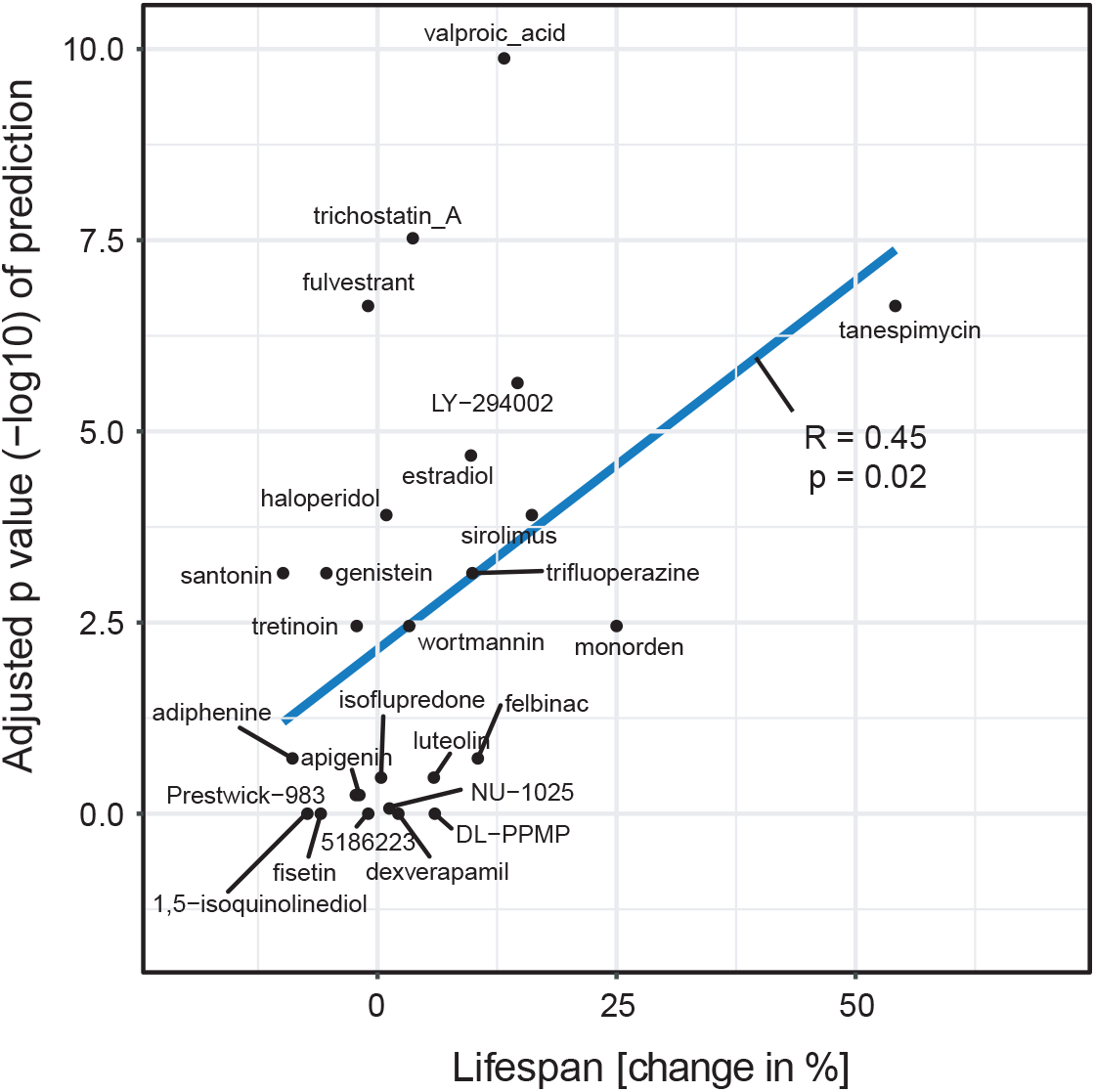
Correlation between our geroprotector predictions and the compounds’ impact on lifespan in *C. elegans*. For the compounds studied in Figure 3, their predicted geroprotective potentials correlate with their ability to extend lifespan in *C. elegans*. The -log10 adjusted p-values of their predictions were plotted against their lifespan phenotypes. The blue line describes a linear regression (R=0.45, p=0.02).

**Figure S5.**
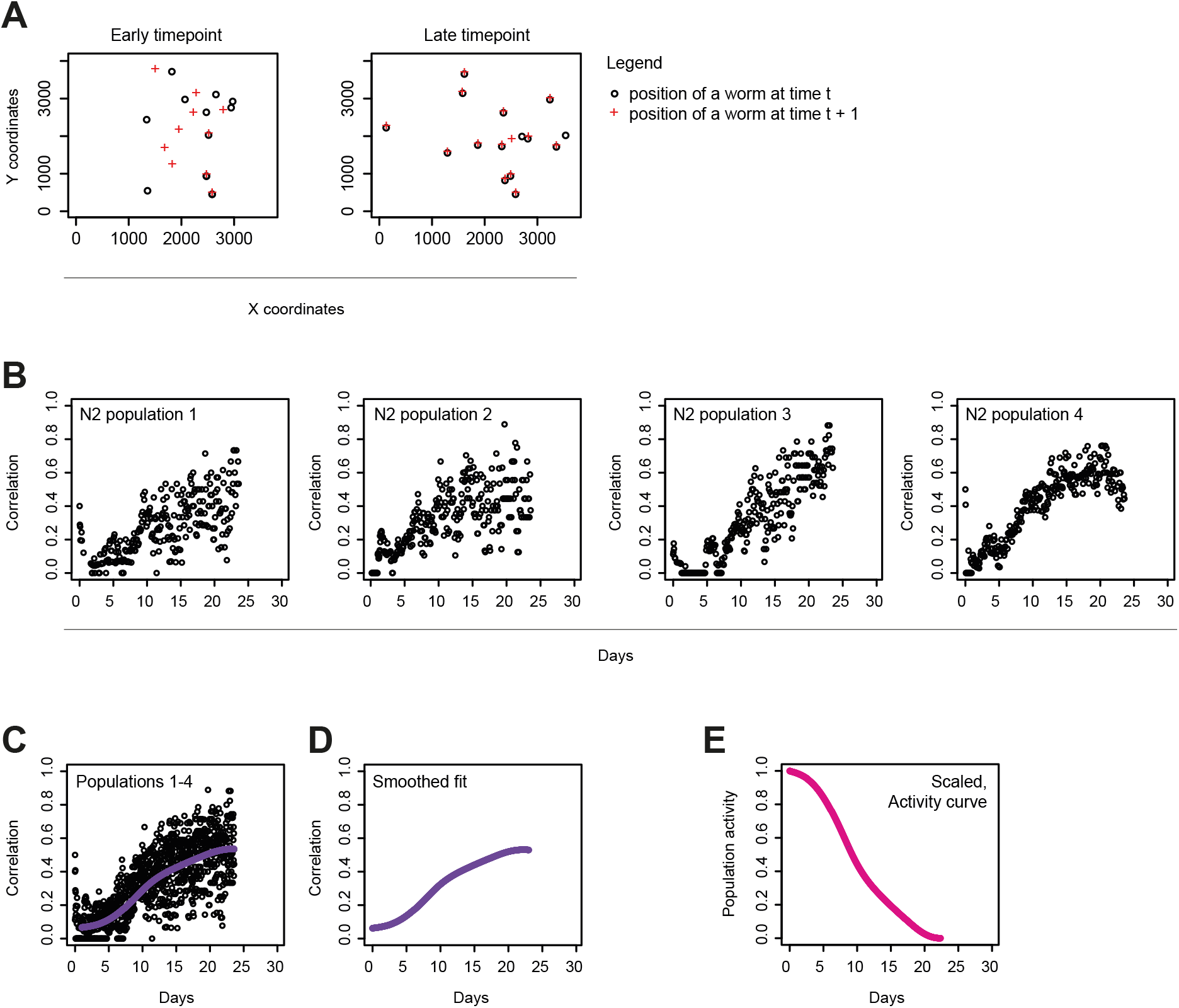
Population health span derived from the ‘lifespan machine’. This is an accompanying figure for steps described in the methods section. (A) Worm positions were assessed based on the position of worm objects identified in scanner images from the ‘lifespan machine’. Left panel: Worm positions identified at an early time point (representing young animals). Abundant changes in worm locations can be seen between consecutive frames. Right panel: The same plot, but at a late time point (representing old animals), showing that worms now remain mostly in the same position. (B) Assessing correlations between subsequent time points as described in (A), shows trends towards increasing correlations in time, indicating less movement of the population. Four independent N2 worm populations are shown. (C) Merging all replicates and fitting a smoothed spline provides a general consensus of the population’s trend. (D) The smoothed spline is considered independently and used to derive the graph shown in (E), by normalizing the starting point to 1 and end point to 0, generating a familiar ‘curve’ that describes the activity of worms at the population level.

**Figure S6.**
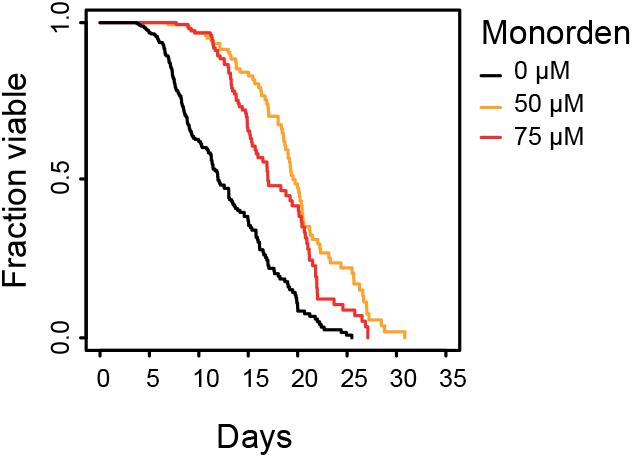
Dose response of Monorden treatment. Survival curve of animals treated either with 50 µM Monorden, 75 μM Monorden, or solvent control, showing that the lifespan benefits of Monorden may already have been maximal at a dose of 50 μM. See table S20 for worm numbers and statistics.

## References

Ayyadevara, S., Engle, M.R., Singh, S.P., Dandapat, A., Lichti, C.F., Beneš, H., Shmookler Reis, R.J., Liebau, E., and Zimniak, P. (2005). Lifespan and stress resistance of Caenorhabditis elegans are increased by expression of glutathione transferases capable of metabolizing the lipid peroxidation product 4-hydroxynonenal. Aging Cell 4, 257–271.

Baird, N. a., Douglas, P.M., Simic, M.S., Grant, A.R., Moresco, J.J., Wolff, S.C., Yates, J.R., Manning, G., and Dillin, A. (2014). HSF-1-mediated cytoskeletal integrity determines thermotolerance and life span. Science 346, 360–363.

Banerji, U. (2009). Heat shock protein 90 as a drug target: some like it hot. Clin. Cancer Res. 15, 9–14.

Bannister, C. a, Holden, S.E., Jenkins-Jones, S., Morgan, C.L., Halcox, J.P., Schernthaner, G., Mukherjee, J., and Currie, C.J. (2014). Can people with type 2 diabetes live longer than those without? A comparison of mortality in people initiated with metformin or sulphonylurea monotherapy and matched, non-diabetic controls. Diabetes. Obes. Metab. 16, 1165–1173.

Barardo, D., Thornton, D., Thoppil, H., Walsh, M., Sharifi, S., Ferreira, S., Anžič, A., Fernandes, M., Monteiro, P., Grum, T., et al. (2017). The DrugAge database of aging-related drugs. Aging Cell 16, 594–597.

Barzilai, N., Crandall, J.P., Kritchevsky, S.B., and Espeland, M.A. (2016). Metformin as a Tool to Target Aging. Cell Metab. 23, 1060–1065.

Bjedov, I., Toivonen, J.M., Kerr, F., Slack, C., Jacobson, J., Foley, A., and Partridge, L. (2010). Mechanisms of life span extension by rapamycin in the fruit fly Drosophila melanogaster. Cell Metab. 11, 35–46.

Blagosklonny, M. V (2002). Hsp-90-associated oncoproteins: multiple targets of geldanamycin and its analogs. Leukemia 16, 455–462.

Calderwood, S.K., Khaleque, M.A., Sawyer, D.B., and Ciocca, D.R. (2006). Heat shock proteins in cancer: Chaperones of tumorigenesis. Trends Biochem. Sci. 31, 164–172.

Calvert, S., Tacutu, R., Sharifi, S., Teixeira, R., Ghosh, P., and de Magalhães, J.P. (2016). A network pharmacology approach reveals new candidate caloric restriction mimetics in C. elegans. Aging Cell 15, 256–266.

Carretero, M., Gomez-Amaro, R.L., and Petrascheck, M. (2015). Pharmacological classes that extend lifespan of Caenorhabditis elegans. Front. Genet. 5.

Chadli, A., Felts, S.J., Wang, Q., Sullivan, W.P., Botuyan, M.V., Fauq, A., Ramirez-Alvarado, M., and Mer, G. (2010). Celastrol inhibits Hsp90 chaperoning of steroid receptors by inducing fibrillization of the co-chaperone p23. J. Biol. Chem. 285, 4224–4231.

Clarke, P. a, Hostein, I., Banerji, U., Stefano, F.D., Maloney, a, Walton, M., Judson, I., and Workman, P. (2000). Gene expression profiling of human colon cancer cells following inhibition of signal transduction by 17-allylamino-17-demethoxygeldanamycin, an inhibitor of the hsp90 molecular chaperone. Oncogene 19, 4125–4133.

Cohen, H.Y. (2004). Calorie Restriction Promotes Mammalian Cell Survival by Inducing the SIRT1 Deacetylase. Science 305, 390–392.

Conte, T.C., Franco, D. V., Baptista, I.L., Bueno, C.R., Selistre-de-Araújo, H.S., Brum, P.C., Moriscot, A.S., and Miyabara, E.H. (2008). Radicicol improves regeneration of skeletal muscle previously damaged by crotoxin in mice. Toxicon 52, 146–155.

Conti, B., Sanchez-Alavez, M., Winsky-Sommerer, R., Morale, M.C., Lucero, J., Brownell, S., Fabre, V., Huitron-Resendiz, S., Henriksen, S., Zorrilla, E.P., et al. (2006). Transgenic mice with a reduced core body temperature have an increased life span. Science 314, 825–828.

Danilov, A., Shaposhnikov, M., Plyusnina, E., Kogan, V., Fedichev, P., and Moskalev, A. (2013). Selective anticancer agents suppress aging in Drosophila. Oncotarget 4, 1507–1526.

David, C.L., Smith, H.E., Raynes, D.A., Pulcini, E.J., and Whitesell, L. (2003). Expression of a unique drug-resistant Hsp90 ortholog by the nematode Caenorhabditis elegans. Cell Stress Chaperones.

Durinck, S., Moreau, Y., Kasprzyk, A., Davis, S., De Moor, B., Brazma, A., and Huber, W. (2005). BioMart and Bioconductor: A powerful link between biological databases and microarray data analysis. Bioinformatics 21, 3439–3440.

Durinck, S., Spellman, P.T., Birney, E., and Huber, W. (2009). Mapping identifiers for the integration of genomic datasets with the R/Bioconductor package biomaRt. Nat. Protoc. 4, 1184–1191.

Echeverría, P.C., Bernthaler, A., Dupuis, P., Mayer, B., and Picard, D. (2011). An interaction network predicted from public data as a discovery tool: application to the Hsp90 molecular chaperone machine. PLoS One 6, e26044.

Evason, K., Collins, J.J., Huang, C., Hughes, S., and Kornfeld, K. (2008). Valproic acid extends Caenorhabditis elegans lifespan. Aging Cell 7, 305–317.

Fridell, Y.W.C., Sánchez-Blanco, A., Silvia, B.A., and Helfand, S.L. (2005). Targeted expression of the human uncoupling protein 2 (hUCP2) to adult neurons extends life span in the fly. Cell Metab. 1, 145–152.

Fuhrmann-Stroissnigg, H., Ling, Y.Y., Zhao, J., McGowan, S.J., Zhu, Y., Brooks, R.W., Grassi, D., Gregg, S.Q., Stripay, J.L., Dorronsoro, A., et al. (2017). Identification of HSP90 inhibitors as a novel class of senolytics. Nat. Commun. 8, 422.

Gentleman, A.R., Carey, V., Huber, W., and Hahne, F. (2017). Genefilter: methods for filtering genes from high-throughput experiments. R Packag. Version 1.58.0.

Griffin, T.M., Valdez, T. V, and Mestril, R. (2004). Radicicol activates heat shock protein expression and cardioprotection in neonatal rat cardiomyocytes. Am. J. Physiol. Heart Circ. Physiol. 287, H1081–H1088.

Gu, Z., and Wang, J. (2013). CePa: An R package for finding significant pathways weighted by multiple network centralities. Bioinformatics 29, 658–660.

Gum, P.A., Thamilarasan, M., Watanabe, J., Blackstone, E.H., and Lauer, M.S. (2001). Aspirin use and all-cause mortality among patients being evaluated for known or suspected coronary artery disease: A propensity analysis. JAMA 286, 1187–1194.

Hahm, J.-H., Kim, S., DiLoreto, R., Shi, C., Lee, S.-J. V., Murphy, C.T., and Nam, H.G. (2015). C. elegans maximum velocity correlates with healthspan and is maintained in worms with an insulin receptor mutation. Nat. Commun. 6, 8919.

Hamilton, B., Dong, Y., Shindo, M., Liu, W., Odell, I., Ruvkun, G., and Lee, S.S. (2005). A systematic RNAi screen for longevity genes in *C. elegans*. Genes Dev. 19, 1544–1555.

Harrison, D.E., Strong, R., Sharp, Z.D., Nelson, J.F., Astle, C.M., Flurkey, K., Nadon, N.L., Wilkinson, J.E., Frenkel, K., Carter, C.S., et al. (2009). Rapamycin fed late in life extends lifespan in genetically heterogeneous mice. Nature 460, 392–395.

Hayes, R.D., Downs, J., Chang, C.K., Jackson, R.G., Shetty, H., Broadbent, M., Hotopf, M., and Stewart, R. (2015). The effect of clozapine on premature mortality: An assessment of clinical monitoring and other potential confounders. Schizophr. Bull. 41, 644–655.

Herranz, D., Muñoz-Martin, M., Cañamero, M., Mulero, F., Martinez-Pastor, B., Fernandez-Capetillo, O., and Serrano, M. (2010). Sirt1 improves healthy ageing and protects from metabolic syndrome-associated cancer. Nat. Commun. 1, 1–8.

Hieronymus, H., Lamb, J., Ross, K.N., Peng, X.P., Clement, C., Rodina, A., Nieto, M., Du, J., Stegmaier, K., Raj, S.M., et al. (2006). Gene expression signature-based chemical genomic prediction identifies a novel class of HSP90 pathway modulators. Cancer Cell 10, 321–330.

Holzenberger, M., Dupont, J., Ducos, B., Leneuve, P., Géloën, A., Even, P.C., Cervera, P., and Le Bouc, Y. (2003). IGF-1 receptor regulates lifespan and resistance to oxidative stress in mice. Nature 421, 182–187.

Hsu, A.-L., Murphy, C.T., and Kenyon, C. (2003). Regulation of aging and age-related disease by DAF-16 and heat-shock factor. Science 300, 1142–1145.

Huang, D.W., Sherman, B.T., and Lempicki, R.A. (2009). Systematic and integrative analysis of large gene lists using DAVID bioinformatics resources. Nat. Protoc. 4, 44–57.

Jung, S.-K., Aleman-Meza, B., Riepe, C., and Zhong, W. (2014). QuantWorm: a comprehensive software package for Caenorhabditis elegans phenotypic assays. PLoS One 9, e84830.

Kaeberlein, M., McVey, M., and Guarente, L. (1999). The SIR2/3/4 complex and SIR2 alone promote longevity in Saccharomyces cerevisiae by two different mechanisms. Genes Dev. 13, 2570–2580.

Kamath, R.S., Fraser, A.G., Dong, Y., Poulin, G., Durbin, R., Gotta, M., Kanapin, A., Le Bot, N., Moreno, S., Sohrmann, M., et al. (2003). Systematic functional analysis of the *Caenorhabditis elegans* genome using RNAi. Nature 421, 231–237.

Kenyon, C., Chang, J., Gensch, E., Rudner, A., and Tabtiang, R. (1993). A C. elegans mutant that lives twice as long as wild type. Nature 366, 461–464.

Kuhn, M., Wing, J., Weston, S., Williams, A., Keefer, C., and Engelhardt, A. (2012). Caret: Classification and Regression Training. Https://Cran.R-Project.Org/Package=Caret.

Kumar, S., and Lombard, D.B. (2016). Finding Ponce de Leon’s Pill: Challenges in Screening for Anti-Aging Molecules. F1000Research 5, 406.

Lamb, J., Crawford, E.D., Peck, D., Modell, J.W., Blat, I.C., Wrobel, M.J., Lerner, J., Brunet, J.-P., Subramanian, A., Ross, K.N., et al. (2006). The Connectivity Map: using gene-expression signatures to connect small molecules, genes, and disease. Science 313, 1929–1935.

Lee, E.B., Ahn, D., Kim, B.J., Lee, S.Y., Seo, H.W., Cha, Y.S., Jeon, H., Eun, J.S., Cha, D.S., and Kim, D.K. (2015). Genistein from vigna angularis extends lifespan in caenorhabditis elegans. Biomol. Ther. 23, 77–83.

Li, Y., Zhang, T., Schwartz, S.J., and Sun, D. (2009). New developments in Hsp90 inhibitors as anti-cancer therapeutics: Mechanisms, clinical perspective and more potential. Drug Resist. Updat. 12, 17–27.

Liaw, a, and Wiener, M. (2002). Classification and Regression by randomForest. R News 2, 18–22.

Liu, J., Lee, J., Hernandez, M.A.S., Mazitschek, R., and Ozcan, U. (2015). Treatment of obesity with celastrol. Cell 161, 999–1011.

Longo, V.D., Antebi, A., Bartke, A., Barzilai, N., Brown-Borg, H.M., Caruso, C., Curiel, T.J., De Cabo, R., Franceschi, C., Gems, D., et al. (2015). Interventions to slow aging in humans: Are we ready? Aging Cell 14, 497–510.

Lonsdale, J., Thomas, J., Salvatore, M., Phillips, R., Lo, E., Shad, S., Hasz, R., Walters, G., Garcia, F., Young, N., et al. (2013). The Genotype-Tissue Expression (GTEx) project. Nat. Genet. 45, 580–585.

Lucanic, M., Garrett, T., Yu, I., Calahorro, F., Asadi Shahmirzadi, A., Miller, A., Gill, M.S., Hughes, R.E., Holden-Dye, L., and Lithgow, G.J. (2016). Chemical activation of a food deprivation signal extends lifespan. Aging Cell 15, 832–841.

Lucanic, M., Plummer, W.T., Chen, E., Harke, J., Foulger, A.C., Onken, B., Coleman-Hulbert, A.L., Dumas, K.J., Guo, S., Johnson, E., et al. (2017). Impact of genetic background and experimental reproducibility on identifying chemical compounds with robust longevity effects. Nat. Commun. 8, 14256.

McCollum, A.K., TenEyck, C.J., Sauer, B.M., Toft, D.O., and Erlichman, C. (2006). Up-regulation of heat shock protein 27 induces resistance to 17-allylamino-demethoxygeldanamycin through a glutathione-mediated mechanism. Cancer Res. 66, 10967–10975.

Mele, M., Ferreira, P.G., Reverter, F., DeLuca, D.S., Monlong, J., Sammeth, M., Young, T.R., Goldmann, J.M., Pervouchine, D.D., Sullivan, T.J., et al. (2015). The human transcriptome across tissues and individuals. Science 348, 660–665.

Mitsui, A., Hamuro, J., Nakamura, H., Kondo, N., Hirabayashi, Y., Ishizaki-Koizumi, S., Hirakawa, T., Inoue, T., and Yodoi, J. (2002). Overexpression of human thioredoxin in transgenic mice controls oxidative stress and life span. Antioxid. Redox Signal. 4, 693–696.

Moskalev, a a, and Shaposhnikov, M. V (2010). Pharmacological inhibition of phosphoinositide 3 and TOR kinases improves survival of Drosophila melanogaster. Rejuvenation Res. 13, 246–247.

Neuwirth, E. (2014). RColorBrewer: ColorBrewer palettes. R Packag. Version 1.1-2 https://cran.Rproject.org/package=RColorBrewer.

Niccoli, T., and Partridge, L. (2012). Ageing as a risk factor for disease. Curr. Biol. 22.

Nussbaum-Krammer, C.I., Neto, M.F., Brielmann, R.M., Pedersen, J.S., and Morimoto, R.I. (2015). Investigating the spreading and toxicity of prion-like proteins using the metazoan model organism C. elegans. J. Vis. Exp. 52321.

Pacey, S., Banerj, U., Judson, I., and Workman, P. (2006). Hsp90 inhibitors in the clinic. Handb. Exp. Pharmacol. 172, 331–358.

Package, T., and Shannon, A.P. (2013). ConnectivityMap: Functional connections between drugs, genes and diseases as revealed by common gene-expression changes. R Packag. Version 1.10.0. 1–3.

Peters, M.J., Joehanes, R., Pilling, L.C., Schurmann, C., Conneely, K.N., Powell, J., Reinmaa, E., Sutphin, G.L., Zhernakova, A., Schramm, K., et al. (2015). The transcriptional landscape of age in human peripheral blood. Nat. Commun. 6, 8570.

Petrascheck, M., Ye, X., and Buck, L.B. (2007). An antidepressant that extends lifespan in adult Caenorhabditis elegans. Nature 450, 553–556.

Porta, C., Paglino, C., and Mosca, A. (2014). Targeting PI3K/Akt/mTOR Signaling in Cancer. Front. Oncol. 4.

Powers, M. V., and Workman, P. (2007). Inhibitors of the heat shock response: Biology and pharmacology. FEBS Lett. 581, 3758–3769.

Pratt, W.B., and Toft, D.O. (2003). Regulation of signaling protein function and trafficking by the hsp90/hsp70-based chaperone machinery. Exp. Biol. Med. (Maywood). 228, 111–133.

Prodromou, C., Roe, S.M., O’Brien, R., Ladbury, J.E., Piper, P.W., and Pearl, L.H. (1997). Identification and structural characterization of the ATP/ADP-binding site in the Hsp90 molecular chaperone. Cell 90, 65–75.

R Core Team (2013). R: A language and environment for statistical computing.

Reboul, J., Vaglio, P., Rual, J.F., Lamesch, P., Martinez, M., Armstrong, C.M., Li, S., Jacotot, L., Bertin, N., Janky, R., et al. (2003). C. elegans ORFeome version 1.1: Experimental verification of the genome annotation and resource for proteomescale protein expression. Nat. Genet. 34, 35–41.

Rogina, B., and Helfand, S.L. (2004). Sir2 mediates longevity in the fly through a pathway related to calorie restriction. Proc. Natl. Acad. Sci. 101, 15998–16003.

Schulte, T.W., Akinaga, S., Soga, S., Sullivan, W., Stensgard, B., Toft, D., and Neckers, L.M. (1998). Antibiotic radicicol binds to the N-terminal domain of Hsp90 and shares important biologic activities with geldanamycin. Cell Stress Chaperones 3, 100–108.

Sharp, S., and Workman, P. (2006). Inhibitors of the HSP90 Molecular Chaperone: Current Status. Adv. Cancer Res. 95, 323–348.

Sidera, K., and Patsavoudi, E. (2014). HSP90 inhibitors: current development and potential in cancer therapy. Recent Pat. Anticancer. Drug Discov. 9, 1–20.

Sing, T., Sander, O., Beerenwinkel, N., and Lengauer, T. (2007). The ROCR Package. R Vignette.

Soga, S., Neckers, L.M., Schulte, T.W., Shiotsu, Y., Akasaka, K., Narumi, H., Agatsuma, T., Ikuina, Y., Murakata, C., Tamaoki, T., et al. (1999). KF25706, a novel oxime derivative of radicicol, exhibits in vivo antitumor activity via selective depletion of Hsp90 binding signaling molecules. Cancer Res. 59, 2931–2938.

Somogyvári, M., Gecse, E., and Sőti, C. (2018). DAF-21/Hsp90 is required for C. elegans longevity by ensuring DAF-16/FOXO isoform A function. Sci. Rep. 8, 12048.

Sonoda, H., Prachasilchai, W., Kondo, H., Yokota-Ikeda, N., Oshikawa, S., Ito, K., and Ikeda, M. (2010). The protective effect of radicicol against renal ischemia--reperfusion injury in mice. J Pharmacol Sci 112, 242–246.

Sood, S., Gallagher, I.J., Lunnon, K., Rullman, E., Keohane, A., Crossland, H., Phillips, B.E., Cederholm, T., Jensen, T., van Loon, L.J.C., et al. (2015). A novel multi-tissue RNA diagnostic of healthy ageing relates to cognitive health status. Genome Biol. 16, 185.

Stiernagle, T. (2006). Maintenance of *C. elegans*. WormBook 1–11.

Stroustrup, N., Ulmschneider, B.E., Nash, Z.M., López-Moyado, I.F., Apfeld, J., and Fontana, W. (2013). The *Caenorhabditis elegans* Lifespan Machine. Nat. Methods 10, 665–670.

Supko, J.G., Hickman, R.L., Grever, M.R., and Malspeis, L. (1995). Preclinical pharmacologic evaluation of geldanamycin as an antitumor agent. Cancer Chemother. Pharmacol. 36, 305–315.

Taipale, M., Jarosz, D.F., and Lindquist, S. (2010). HSP90 at the hub of protein homeostasis: emerging mechanistic insights. Nat. Rev. Mol. Cell Biol. 11, 515–528.

Tao, D., Lu, J., Sun, H., Zhao, Y.-M., Yuan, Z.-G., Li, X.-X., and Huang, B.-Q. (2004). Trichostatin A extends the lifespan of Drosophila melanogaster by elevating hsp22 expression. Acta Biochim. Biophys. Sin. (Shanghai). 36, 618–622.

Taskesen, E., and Reinders, M.J.T. (2016). 2D representation of transcriptomes by t-SNE exposes relatedness between human tissues. PLoS One 11, 1–6.

Tatar, M., Kopelman, A., Epstein, D., Tu, M.P., Yin, C.M., and Garofalo, R.S. (2001). A mutant Drosophila insulin receptor homolog that extends life-span and impairs neuroendocrine function. Science 292, 107–110.

The GTEx Consortium (2015). The Genotype-Tissue Expression (GTEx) pilot analysis: Multitissue gene regulation in humans. Science 348, 648–660.

Therneau, T. (2012). A Package for Survival Analysis in S. R package version. Survival (Lond).

Tissenbaum, H. a, and Guarente, L. (2001). Increased dosage of a sir-2 gene extends lifespan in Caenorhabditis elegans.

Trepel, J., Mollapour, M., Giaccone, G., and Neckers, L. (2010). Targeting the dynamic HSP90 complex in cancer. Nat. Rev. Cancer 10, 537–549.

Trott, A., West, J.D., Klaic, L., Westerheide, S.D., Silverman, R.B., Morimoto, R.I., and Morano, K.A. (2007). Activation of Heat Shock and Antioxidant Responses by the Natural Product Celastrol: Transcriptional Signatures of a Thiol-targeted Molecule. Mol. Biol. Cell 19, 1104–1112.

Tukaj, S., and Węgrzyn, G. (2016). Anti-Hsp90 therapy in autoimmune and inflammatory diseases: a review of preclinical studies. Cell Stress Chaperones 21, 213–218.

Umeda-Kameyama, Y., Tsuda, M., Ohkura, C., Matsuo, T., Namba, Y., Ohuchi, Y., and Aigaki, T. (2007). Thioredoxin suppresses parkin-associated endothelin receptor-like receptor-induced neurotoxicity and extends longevity in Drosophila. J. Biol. Chem. 282, 11180–11187.

Wasko, M.C.M., Dasgupta, A., Hubert, H., Fries, J.F., and Ward, M.M. (2013). Propensity-adjusted association of methotrexate with overall survival in rheumatoid arthritis. Arthritis Rheum. 65, 334–342.

Welch, W.J., and Feramisco, J.R. (1982). Purification of the major mammalian heat shock proteins. J. Biol. Chem. 257, 14949–14959.

Welch, W.J., Kang, H.S., Beckmann, R.P., and Mizzen, L.A. (1991). Response of mammalian cells to metabolic stress; changes in cell physiology and structure/function of stress proteins. Curr. Top. Microbiol. Immunol. 167, 31–55.

Westerheide, S.D., Bosman, J.D., Mbadugha, B.N.A., Kawahara, T.L.A., Matsumoto, G., Kim, S., Gu, W., Devlin, J.P., Silverman, R.B., and Morimoto, R.I. (2004). Celastrols as inducers of the heat shock response and cytoprotection. J. Biol. Chem. 279, 56053–56060.

Whitesell, L., and Lindquist, S.L. (2005). HSP90 and the chaperoning of cancer. Nat. Rev. Cancer 5, 761–772.

Wu, C. (1995). Heat shock transcription factors: structure and regulation. Annu. Rev. Cell Dev. Biol. 11, 441–469.

Yang, J., Huang, T., Petralia, F., Long, Q., Zhang, B., Argmann, C., Zhao, Y., Mobbs, C. V., GTEx Consortium, Schadt, E.E., et al. (2015). Synchronized age-related gene expression changes across multiple tissues in human and the link to complex diseases. Sci. Rep. 5, 15145.

Ye, X., Linton, J.M., Schork, N.J., Buck, L.B., and Petrascheck, M. (2014). A pharmacological network for lifespan extension in Caenorhabditis elegans. Aging Cell 13, 206–215.

Zarse, K., and Ristow, M. (2008). Antidepressants of the serotonin-antagonist type increase body fat and decrease lifespan of adult Caenorhabditis elegans. PLoS One 3.

Zhao, Y., Huang, Z.J., Rahman, M., Luo, Q., and Thorlacius, H. (2013). Radicicol, an Hsp90 inhibitor, inhibits intestinal inflammation and leakage in abdominal sepsis. J. Surg. Res. 182, 312–318.

Zou, J., Guo, Y., Guettouche, T., Smith, D.F., and Voellmy, R. (1998). Repression of heat shock transcription factor HSF1 activation by HSP90 (HSP90 complex) that forms a stress-sensitive complex with HSF1. Cell 94, 471–480.

